# Structural Mechanism of Filamentation Induced Dampening of GTP Inhibition of Glutamate Dehydrogenase

**DOI:** 10.64898/2026.07.06.736867

**Authors:** Zelin Shan, Noura I. Darwish, Andres Rivero-Gamez, Timothy C. Strutzenberg, Dmitry Lyumkis, Nancy C. Horton

## Abstract

Glutamate dehydrogenase (GDH) is a highly regulated key enzyme that catalyzes the reversible oxidative deamination of glutamate to α-ketoglutarate, positioning it at a critical hub linking amino acid catabolism to energy production while supplying ammonia for urea synthesis and other nitrogen pathways. Early investigations have shown that bovine GDH (bGDH), which shares 98% sequence identity with its human homolog, assembles into polymeric filaments with altered allosteric responses. Filamentation has only relatively recently been appreciated as a widespread mechanism of enzyme regulation, prompting a reevaluation of these early observations in GDH. Here, we use high-resolution cryogenic electron microscopy (cryo-EM) to show that bGDH hexamers assemble via reciprocal “antenna” interactions that oppose the conformational changes associated with GTP inhibition, revealing how filamentation reshapes GDH allostery and with implications for the treatment of human disease.

## INTRODUCTION

Glutamate dehydrogenase (GDH, EC 1.4.1.3) is a conserved metabolic enzyme that catalyzes the reversible oxidative deamination of L-glutamate to -ketoglutarate, ammonia, and NAD(P)H, linking amino acid metabolism to the tricarboxylic acid (TCA) cycle, nitrogen metabolism, and cellular redox balance^1^. In humans, two mitochondrial isoforms are encoded by distinct genes, GLUD1 and GLUD2. GLUD1 (GDH1) is a broadly expressed “housekeeping” enzyme with particularly high levels in liver, pancreas, kidney, and brain. GLUD2 (GDH2) is a primate-specific isoform whose expression is largely confined to neural, testicular, and other steroidogenic or endocrine tissues (including astrocytes and subsets of cortical neurons)^2–4^. The reaction catalyzed by GDH is readily reversible *in vitro* and near equilibrium thermodynamically, but *in vivo* its net flux is usually biased toward oxidative deamination or, less commonly, reductive amination, depending on substrate/product concentrations, cofactor ratios, and allosteric regulation^1^. In most mammalian tissues, GDH1 is thought to support oxidative deamination of glutamate (Glu) to supply -ketoglutarate for the TCA cycle, thereby coupling amino acid catabolism to ATP production. In liver and kidney, this activity also contributes to gluconeogenesis and to ammonia handling for urea synthesis^1^.

Given its significant role in mitochondrial bioenergetics and amino acid metabolism, human GDH1 (hGDH1) is critical to the metabolic function of numerous cell types and tissues. hGDH1 dysfunction is best characterized in patients with hyperinsulinism/hyperammonemia (HI/HA) syndrome, where hGDH1 mutations cause inappropriate, amino acid–stimulated insulin secretion and chronically elevated serum ammonium levels^1^. GDH1 also coordinates neurotransmitter and amino acid homeostasis within the brain^5–8^, and partial hGDH1 deficiency has been reported in patients with cerebellar degeneration^9^, underscoring its importance for neuronal function. More broadly, altered hGDH1 activity supports metabolic reprogramming in complex diseases such as cancer and diabetes, highlighting its potential as a therapeutic target^1,2^.

The formation of filaments, or linear/helical polymers, by enzymes from a wide distribution of biochemical pathways is now recognized as an important layer in enzyme regulation^10–12^. Early studies with bovine GDH (bGDH), which shares 98% sequence identity with its human homolog hGDH1, demonstrated that the enzyme forms polymeric filaments *in vitro*^13–15^. For example, the first reported measurement of bGDH molecular weight was conducted by Olson and Anfinsen in 1952 using sedimentation analysis indicating that the native form of bGDH is approximately 1 MDa and composed of multiple associated species^16^, which are now known to be bGDH hexamers with a molecular weight of approximately 340 kDa each. Related studies found that bGDH could independently associate into higher-order arrangements, such as linear polymers, multistrand rods, and sheets^14,15,17–22^. The formation and stabilization of bGDH filaments appears to be dependent on protein concentration, substrate availability, solution conditions, and the presence or absence of allosteric effectors^23,24^. GDH is highly enriched in liver mitochondria and estimates place its matrix concentration in the low mg/ml range^25,26^, a regime that *in vitro* corresponds predominantly to filamentous assemblies^15,17,23,24^.

The forward oxidative deamination reaction catalyzed by GDH involves multiple distinct steps: NAD(P)^+^ and Glu bind to an open enzyme conformation which then triggers active site closing allowing the reaction to proceed. The active site then re-opens to release the products NH_3_, NAD(P)H, and αKG^27^. However, this simple reaction pathway is regulated through complex processes. For example, physiological Glu concentrations and pH lead to the formation of an abortive complex, where Glu rebinds in the active site after release of αKG and NH_3_ but before the release of the reduced coenzyme NAD(P)H, thereby inhibiting further catalysis^1,19^.

Several known allosteric effectors control bGDH activity by controlling the rate of dissociation of this abortive complex^1,19,28^. GTP stabilizes the closed active site conformation, thereby preventing abortive complex dissociation and inhibiting the enzyme. ADP reverses the effect of GTP by allowing active site opening^29,30^. Interestingly, NADH (but not NADPH) also binds in the ADP binding site, particularly in the presence of GTP and Glu, to stabilize the inhibited closed conformation^1,29,30^. Hence, GTP, ADP, and NADH act as strong allosteric regulators of bGDH activity by controlling enzyme conformation, and in particular active site opening. bGDH is also regulated by several other allosteric effectors, such as leucine, palmitoyl-CoA, and certain steroid hormones^1^.

Several classic studies have shown that filamentation of bGDH strongly influences how it responds to GTP and NADH. For example, filamentation of bGDH reduces its sensitivity to inhibition by guanosine nucleotides (GTP or GDP) as well as by NADH^24,31^. Further, in the presence of reduced coenzyme, GTP binds preferentially to hexameric (i.e. non-filamented) bGDH, but can also bind to filamentous bGDH, albeit with lower affinity, and drive filament dissociation to the hexameric form^24,32–34^. These observations indicate that filamentation attenuates allosteric inhibition by GTP and NADH, however the underlying structural mechanism had remained undefined until now.

To define the molecular basis for how filamentation alters the allosteric response of GDH, we determined four new high-resolution structures of bGDH, representing filamentous and non-filamentous states, and in apo and liganded forms. The structures reveal the mechanism by which discrete bGDH hexamers engage one another to regulate bGDH filamentation and function. We propose a model wherein interactions between bGDH hexamers modulate their response to allosteric effectors by opposing the conformational changes induced by formation of the GTP (and NADH) stabilized abortive complex.

## RESULTS

### bGDH forms filaments in a concentration dependent manner

bGDH resides in a dynamic equilibrium between monomers, dimers, trimers, hexamers, and filamentous assemblies^35^, with the hexamer acting as the minimal building block of the higher-order filaments^15,36^. We confirmed the self-assembly of bGDH in a buffer mimicking physiological conditions using mass photometry, electron microscopy, and analytical ultracentrifugation and with bGDH concentrations ranging from 1.25 μg/ml to 0.5 mg/ml (**Fig. S1, Tables S1-S3**). These particular techniques were chosen to best analyze low, intermediate, and high enzyme concentrations, respectively, for the formation of enzyme filaments. At low concentrations (1.25 - 10 μg/ml), most bGDH is found in the hexameric form, with fewer than 5% of hexamers self-associating into larger assemblies (**Fig. S1a**, **Table S1**). At intermediate concentrations (50 μg/ml), ∼60% of bGDH forms short filaments containing 2-4 hexamers (**Fig. S1b**, **Table S2**). At the highest concentrations (0.5-1.5 mg/ml), ∼60% of bGDH was found in self-assemblies ranging from three to seven associated hexamers (**Table S3**). However, a significant amount of enzyme was also found to precipitate during centrifugation at these high bGDH concentrations, leading to an underestimation of the degree of self-assembly formation. The sedimentation analysis was also performed at three different speeds, and the higher speeds introduced a concentrating effect in the sample that increased the proportion of high molecular weight species (HWMS, defined as species that include at least two hexamers)(**Table S3**). Together, the data from the three methods plotted in **Figure S1d** shows that the percentage of HMWS increases as a function of bGDH concentration, reaching at least ∼60% of total enzyme concentration at 0.5 mg/ml bGDH.

### Dynamic equilibrium of bGDH hexamers and filament stabilization

Although bGDH forms filamentous higher order assemblies under physiological conditions, high concentrations of ammonium sulfate or sodium sulfate^15^ as well as chemical crosslinking^21,31,37^ are known to stabilize the bGDH filaments. At low ionic strength and low concentrations of bGDH, we predominantly observe monodisperse bGDH hexamers, with a few short linear filaments (**Fig. S2a**). Chemical crosslinking prevents the dissociation of hexamers from filaments, resulting in longer and more abundant filaments under otherwise identical buffer conditions (**Fig. S2b**). The self-assemblies are regular and roughly linear, consistent with filamentation rather than aggregation. The ability to trap these species with chemical crosslinking indicates that hexamer-hexamer associations occur readily under physiological conditions but also occur transiently. As previously reported^15^, filaments assembled in the presence of buffer containing 1 M sodium sulfate were longer and more abundant (**Fig. S2c**). The presence of both 1M sodium sulfate and glutaraldehyde crosslinker yielded the longest and most densely populated filaments on negatively stained grids (**Fig. S2d**). Notably, under all tested conditions, bGDH filaments showed the same overall morphology as observed in the absence of any treatment, indicating that chemical treatments help stabilize filamentous assemblies but do not fundamentally alter the interactions between hexamers.

To aid structural studies, we treated bGDH samples with crosslinker in the presence of 600 mM or 1 M sodium sulfate and isolated polymeric assemblies via size-exclusion chromatography (SEC). We selected a leading segment of the broad peak for subsequent grid preparation (**Fig. S3a-b**). These procedures yielded populations of bGDH that were enriched in larger bGDH assemblies for both apo and liganded samples, facilitating downstream structural analysis.

### Symmetric interactions utilizing the antenna drive bGDH filamentation

To gain insights into the molecular basis by which filamentation modulates bGDH structure and function, we set out to determine high-resolution structures of bGDH assemblies in non-filamentous and filamentous states. We initially focused on the apo form and collected 3,480 movies of frozen, hydrated apo bGDH complexes that had been crosslinked and purified via SEC. 2D class averages revealed the presence of both monomers and multimers of hexameric bGDH assemblies (**Fig. S4)**. Individual non-filamented bGDH hexamers (i.e. the mono-hexamer) refined to 2.5 Å resolution using single-particle processing protocols in cryoSPARC^38^, while the combination of density subtraction and local refinement also yielded a map of a dimer of bGDH hexamers (the di-hexamer). For the di-hexamer, the primary hexamer and the neighboring hexamer were resolved to 2.6 Å and 2.9 Å, respectively. We derived atomic models for all maps (**Figs. S4-S6, Table S4-S5)**. Because the di-hexameric form is the smallest multimeric component of filamentous assemblies, we subsequently refer to this as the minimal filamentous state.

The mono-hexameric bGDH configuration in its apo form consists of six identical protomers arranged within a configuration defined by D3 symmetry (**Fig. S6a-b**). This organization is well-known from previously determined crystal structures^29,30,39^. As seen in the previously determined structures of mono-hexameric bGDH, pyramidal structures known as “antenna” project from either end along the three-fold symmetry axis of the hexamer, which are formed by the assembly of a long antenna helix from each of three subunits. In the di-hexameric configuration, bGDH hexamers self-associate at two contact points in a two-fold symmetric manner. Each contact point involves a region known as the NAD binding domain of one subunit within one hexamer and the tip of an antenna of the other hexamer (**Fig. 1a**). A view down the 3-fold axis of the mono- and di-hexamer antenna structure is shown in **Fig. 1b-c**. At the interface between hexamers, the side chain of Pro240 (or Pro297 in the unprocessed protein) of hexamer_1_ rests inside a hydrophobic cup at the antenna tip created by a trio of side chains Phe421 (or Phe478 in the unprocessed form) from each chain of hexamer_2_ (**Fig. 1d**). This interaction breaks the internal symmetry in the antenna, producing a large outward movement of the loops adjacent to Phe421 (**Fig. 1e, Movie S1)**. Loop displacement is not symmetric, ranging from a root mean square deviation (RMSD) of 5.4 Å to 6.2 Å in the three chains, quantified between residues 423-425, when compared to the mono-hexamer structure. Hence, the hexamer–hexamer interface is symmetry-mismatched: the threefold-symmetric tip of one hexamer contacts only a single subunit of the neighboring hexamer, resulting in the adoption of distinct conformations in the three subunits at the antenna tips. This kind of local symmetry breaking in otherwise symmetric assemblies has been noted in other protein complexes^40^.

**Figure 1.**
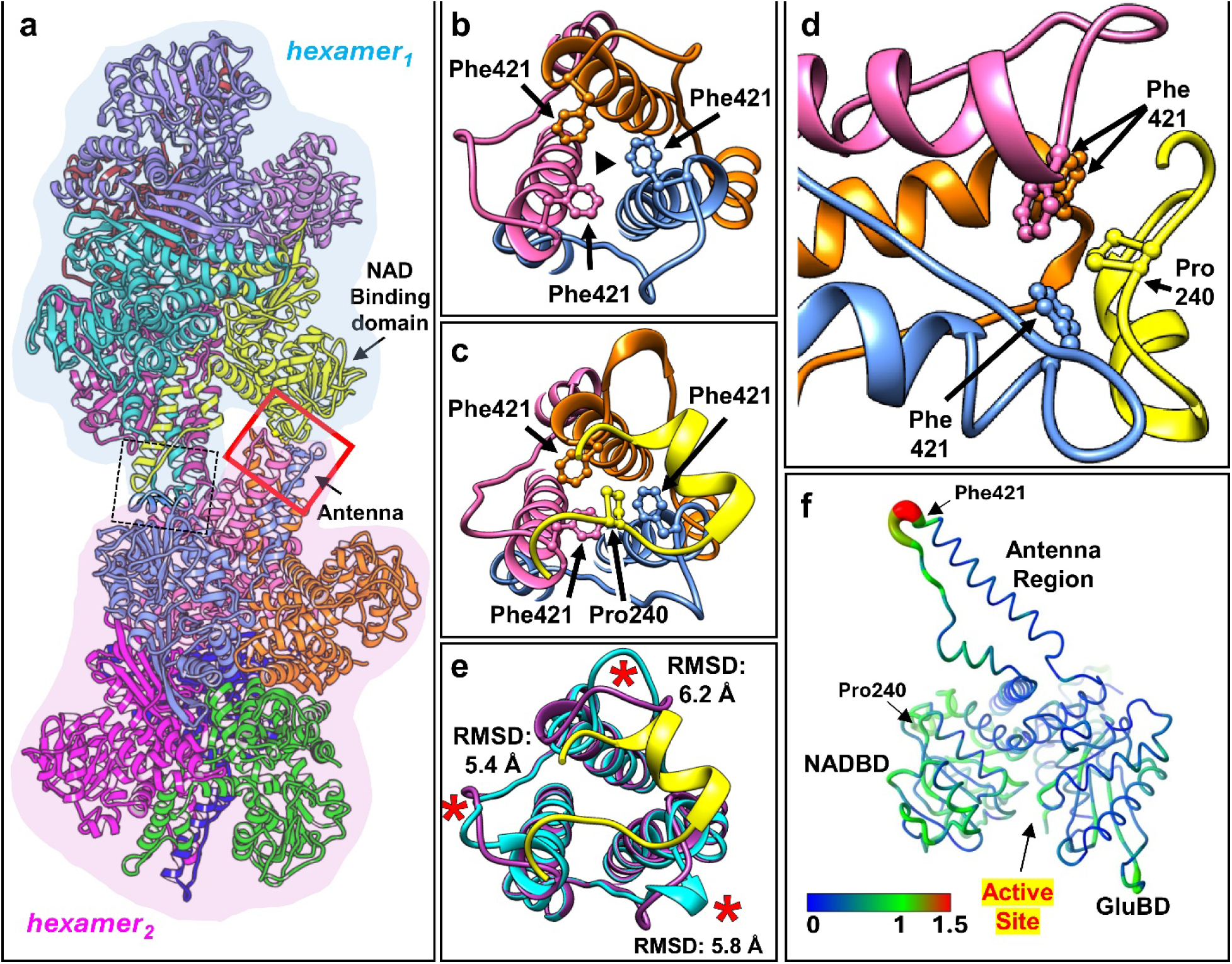
Analysis of bGDH in mono- and di-hexamer structures in the apo (unliganded) state. **a,** Ribbon diagram of the di-hexamer structure in the apo state. The region boxed in red identifies one of two symmetric contacts (the other outlined by a dotted black box) between the hexamers involving the antenna of hexamer_2_ (orange, pink, blue) and a subunit of hexamer_1_ (yellow). **b,** Close-up view down the 3-fold axis (black triangle) of an antenna of the mono-hexamer. The Phe421 side chain in each of three chains is shown in ball and stick. **c,** Close-up view down the 3-fold axis of the contact between hexamers boxed in (a). Phe421 from the antenna (pink, blue, orange) and Pro240 from of the neighboring hexamer (yellow) are shown in ball-and-stick. The interaction between hexamers breaks the 3-fold symmetry in the antenna. **d,** Side view of the contact shown in (c). **e,** Overlay of antenna residues from mono-(dark purple) and di-hexamer (cyan, yellow) structures. Asterisks indicate residues 423-425, which show high RMSD between the two structures. **f,** A single subunit from the apo mono-hexamer, with the position of the antenna region, NAD binding domain (NADBD) and glutamate binding domain (GluBD) indicated. The active site resides between the NADBD and GluBD. RMSD values from the comparison of a single subunit between the apo mono- and di-hexamer structures are mapped onto the subunit. RMSD values (Å) are shown by color and chain width, with thicker chains corresponding to higher RMSD. The Cα atoms of the GluBD residues (residues 5-208, 474-496, 369-390) were used for the superposition. The highest RMSD occur in the antenna helix and the adjacent descending loop.

The apo bGDH subunits in both mono- and di-hexamer forms reside in the open active site conformation, typical of ligand-free bGDH^30^. Alignment of an individual protomer from the mono- and di-hexameric reconstructions indicates that no other significant conformational changes arise in regions outside the antenna, including the functionally relevant NAD binding domain (NADBD) containing Pro240, and the glutamate binding domain (GluBD) (**Fig. 1f**). In addition, the interface between interacting hexamers does not overlap or closely approach any other known ligand binding site. Thus, the conformations of the ligand binding sites and the active sites are not altered by bGDH filamentation, with both the non-filamenting and filamenting forms preserving the open conformation in all six protomers. Taken together, the structures indicate that symmetric interactions via hexamer antennae stabilize bGDH filamentation without obstructing access of ligands or substrates to their binding sites.

**Figure S7** shows models of longer filaments built from the di-hexamer coordinates. Because of the three-fold symmetry of the hexamer at the antenna, and the asymmetry between interacting hexamers (only one subunit of each bGDH hexamer engages at the interface with its NAD binding domain), the addition of each bGDH hexamer (of a di-hexamer unit) to the growing filament can take on one of three possible orientations. One orientation leads to a straight filament (**Fig. S7a**), a second leads to a filament with an approximate 30° bend between di-hexamers (**Fig. S7b**), and a third produces a filament with a left-handed twist and an ∼60° bend between adjacent di-hexamers (**Fig. S7c**). Of course, filaments need not contain purely one type of contact and can contain mixtures of the three leading to more irregular appearing filaments, as seen in images (e.g. **Fig. S2**).

### Liganded state structures of non-filamentous and filamentous bGDH

In order to better understand how filamentation affects the response of bGDH to GTP, we determined the structures of bGDH with GTP, Glu, and NADH in both mono- and di-hexameric forms. This combination of ligands bound to bGDH results in the structurally well characterized closed active site state^29^. The binding of Glu and NADH within the active site of bGDH results in a complex known as the inhibited “abortive complex”, which is stabilized by the binding of GTP (**Fig. 2a**). In addition, at sufficient concentrations, a second molecule of NADH will bind to a separate allosteric effector site within each bGDH subunit, further stabilizing the abortive complex^29^. Using similar procedures as those described above, the high-resolution structure of mono-hexameric bGDH was resolved to 2.2 Å, and the di-hexameric bGDH structure was resolved to 3.3 Å and 3.5 Å for the two hexamers, respectively, with corresponding refined atomic models (**Figs. S8-S10**)(**Tables S4-S5**).

**Figure 2.**
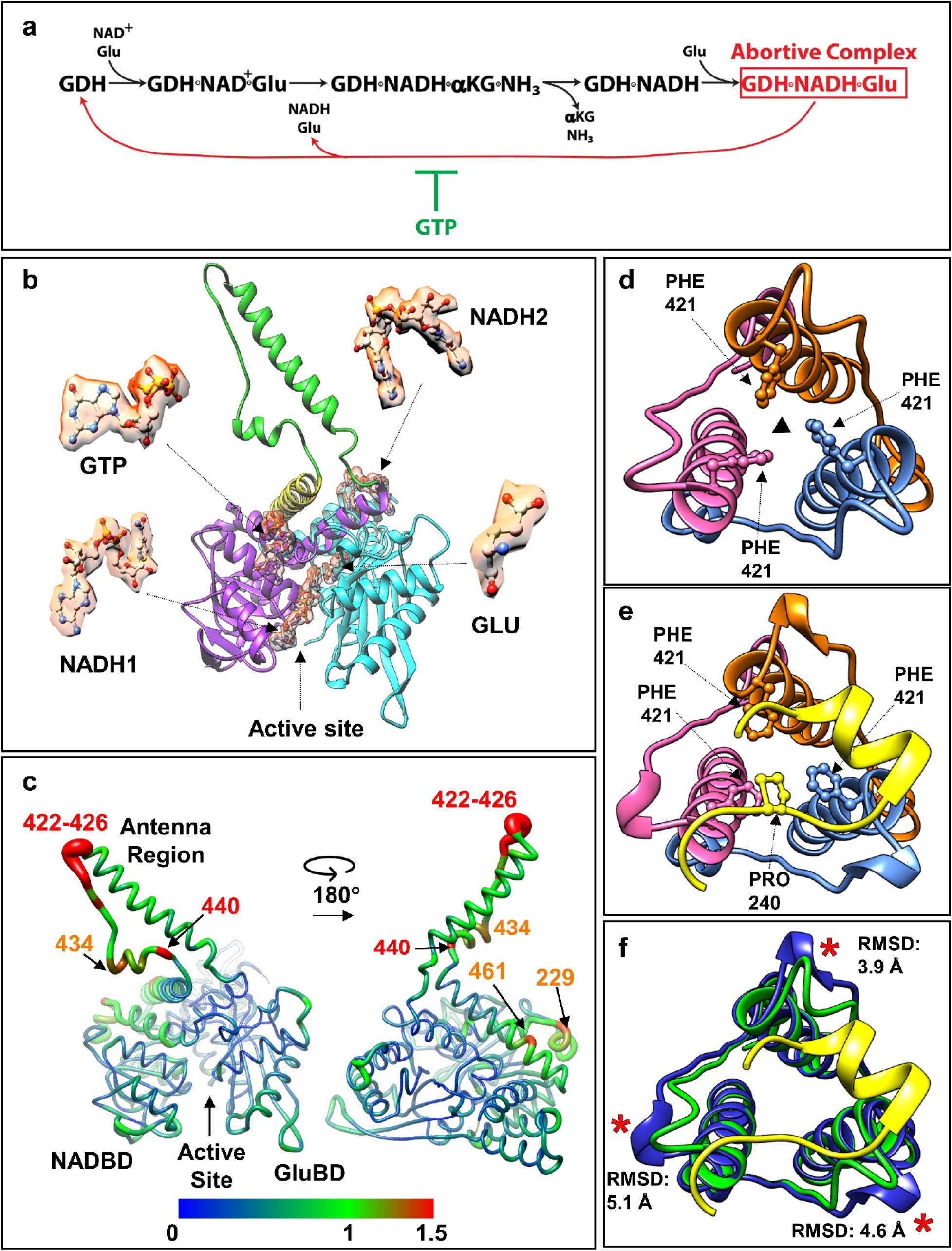
Analysis of liganded mono- and di-hexameric structures of bGDH. **a,** GDH binds converts NAD^+^ (or NADP+) and glutamate (Glu) to NADH (or NADPH), alpha-ketoglutarate (αKG), and ammonia (as well as one proton, not shown). The products αKG and ammonia are released. A second molecule of Glu binds before the reduced coenzyme NADH (or NADPH) is released thereby forming the abortive, inhibited complex. GTP binds in an allosteric site and stabilizes the abortive complex. Sufficient concentrations of NADH (but not NADPH) result in its binding at a second distinct allosteric site which further inhibits bGDH (not shown). **b,** A single subunit of bGDH from the liganded mono-hexameric structure showing the ligand binding sites and their corresponding density in cryo-EM maps (antenna region, green; NADBD, pink; GluBD, cyan). **c,** RMSD (Å) between mono- and di-hexameric bGDH subunits in the liganded state. RMSD values (Å) are shown by color and chain width, with thicker chains corresponding to higher RMSD. The Cα atoms of the GluBD (residues 5-208, 474-496, 369-390) were used for superposition. The subunit of the di-hexamer involved in hexamer-hexamer contacts via its NAD binding domain, NADBD, and antenna helix, was used for the superposition. The highest RMSDs occur in the long antenna helix, the adjacent descending loop, and the short helix containing residues Ala434 and Ile440. **d,** Close-up view down the 3-fold axis (black triangle) of an antenna domain of the liganded, mono-hexameric bGDH structure. Phe421 from each chain is shown in ball-and-stick. **e,** As in (c), but for the di-hexamer structure of liganded bGDH. A segment of the NAD binding domain of the contacted neighboring hexamer is shown in yellow, and residue Pro240 in ball-and-stick. **f,** Close-up view down the 3-fold axis of the antenna of the liganded bGDH mono-hexamer (green) and di-hexamer (blue, yellow) structures. The asterisk indicates the position of residues 422-426, which show high RMSDs between the liganded mono- and di-hexamer structures.

In both the liganded mono- and di-hexamer structures, one Glu and one NADH bind in each active site of all six subunits, similar to previously determined crystal structures of hexameric bGDH^29^. GTP binds in the allosteric GTP binding site, between the antenna and NAD binding domain of each subunit, and a second copy of NADH is bound in the ADP binding site of each subunit (**Fig. 2b**). Concomitant with ligand binding, a large conformational change shifts the NAD binding domain towards the Glu binding domain in each subunit (**Movie S2**). This reorientation also involves a shift in the position of the antenna, and a change in conformation in the small helix found at the base of the antenna (see **Fig. S11** for RMSD analysis and **Fig. S12a-b** for superpositions of apo and liganded mono-hexamer structures). Overall, the structure is similar to x-ray crystal structures of non-filamentous bGDH containing the same bound ligands, constituting the abortive, inhibited, closed complex^29^.

With ligand binding, structural differences are more substantial between mono- and di-hexamers than in the apo structure. Higher RMSDs characterize the tips of the antenna helices spanning residues 422-426, as well as the short helix at 434–440, and the surface of the NAD binding domain around residues 229 and 461 (**Fig. 2c**). The differences are particularly pronounced within the subunits located at the hexamer-hexamer interface, which involve their NAD binding domains (**Fig. S12c-d, Movie S3**). Compared to the apo mono-hexamer structure, the antenna tip of the liganded mono-hexamer structure shows a similar, albeit non-identical conformation (**Fig. 2d**). In the liganded di-hexamer structure, Pro240 of one hexamer is again positioned in the hydrophobic cup created from the three antenna helices containing Phe421 from the interacting hexamer (**Fig. 2e**). However, the position of the Pro240 segment is shifted in this interface compared to that found the apo di-hexamer structure (**Movie S4**). Furthermore, the relative subunit arrangement in the liganded di-hexamer has changed when compared with the apo form, such that there are now more pronounced differences between the mono- and di-hexamer within individual subunits. (**Fig. S13**). Finally, local distortions resulting from the hexamer-hexamer interaction are found in the antenna tip, similar but not identical to those found in the apo di-hexamer structure (**Fig. 2f**). These differences and their implications will be discussed below.

### The hexamer-hexamer contact is more dynamic in the liganded di-hexamer structure

Ligand binding and closing of the active site via a shift in the position of the NAD binding domain results in a shift in the position of the Pro240 loop of hexamer_1_ relative to the interacting antenna domain of hexamer_2_ (**Fig. 3a-b**). In addition, filamentous assemblies of liganded hexamers in negative stain micrographs were less populous on grids, shorter, and characterized by increased flexibility at the hexamer-hexamer interface in comparison to the apo assemblies (**Fig. S14**). 2D class averages from the negatively stained data also showed that at most two bGDH hexamers were averageable in the liganded form, whereas three hexamers were averageable in the apo forms; the second hexamer was also typically less featureful in the liganded dataset (**Fig. 3c-d**). In high-resolution cryo-EM experiments, the liganded di-hexamers were consistently resolved to a lower resolution than the apo di-hexamers, despite more particles contributing to reconstructions (**Figs. S4-S6** and **Figs. S8-S10**).

**Figure 3.**
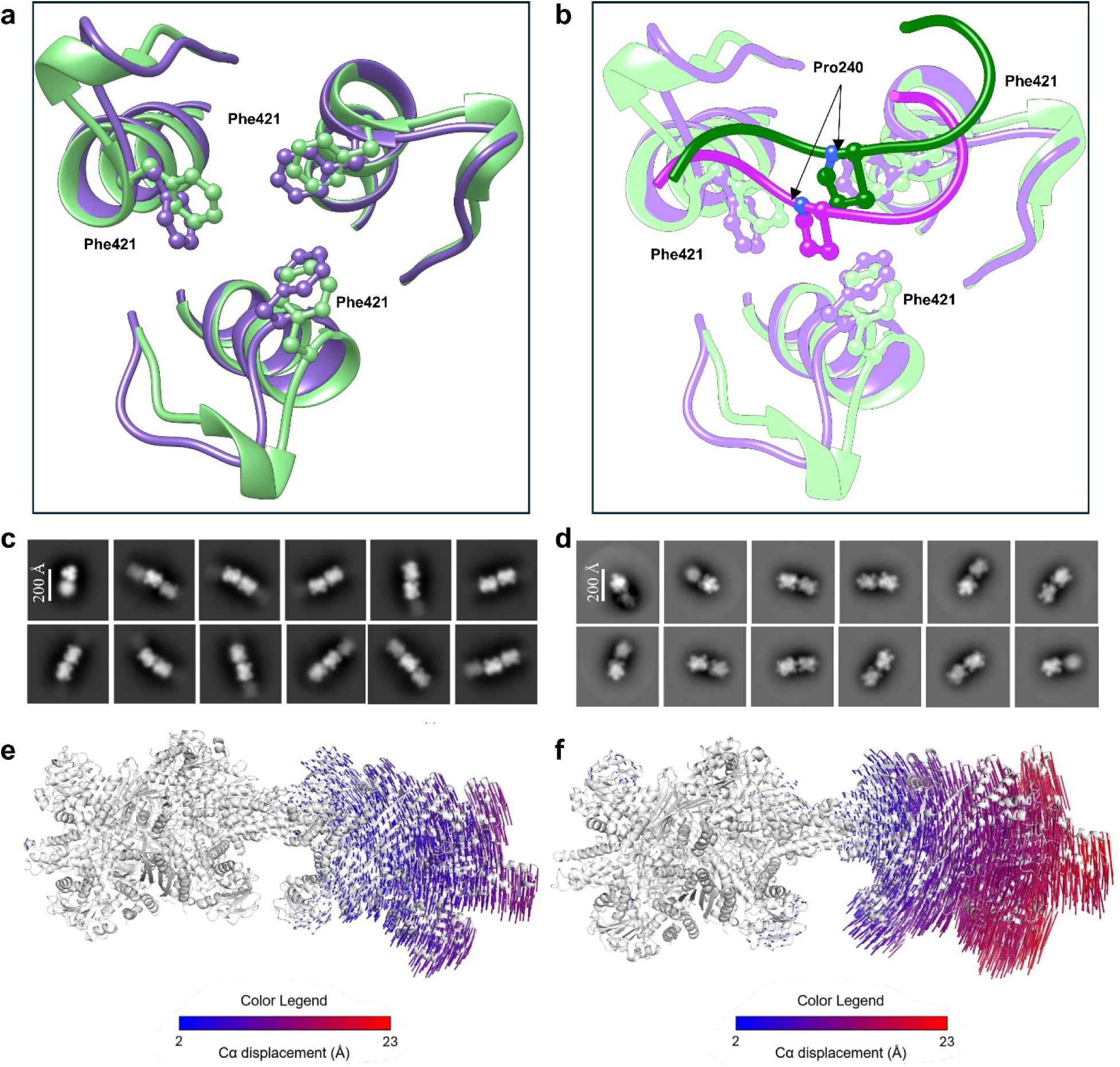
The hexamer-hexamer interface changes with ligand binding. **a,** Superposition of apo (dark purple) and liganded (green) di-hexamer structures using the Cα of residues 398-420 of the antenna helix from one subunit. The side chains of Phe421 from each subunit are shown in ball and stick. **b,** As in (a), now including the Pro240 loop from the contacting hexamer of the apo (plum) and liganded (green) di-hexamer structures. In the liganded structure, the position of Pro240 is shifted relative to the antenna residues. **c-d,** Representative 2D class averages from negatively stained images show filamentous bGDH in (c) apo and (d) liganded states. Scale bar is indicated. **e-f,** 3DFlex analyses showing the variability of the hexamer-hexamer interface in the (**e**) apo and (**f**) liganded states. Variability is quantified by Cα displacement, represented by the color legend and vector length.

To quantify the conformational dynamics of bGDH filamentation and define the movement about the hexamer-hexamer interface, we used 3D flexibility analysis^41^. We trained a 3DFlex model on the dimer-of-hexamer particle stacks from the apo and liganded cryo-EM datasets and used the model to compute learned ranges of motion (**Movies S5-S6**). We relaxed atomic models into the maps, aligned the ensemble of models to a single subunit, and calculated global backbone RMSD values and dihedral bend angles for the neighboring hexamer. The apo and liganded forms are characterized, respectively, by ∼1.5 Å and ∼3.0 Å RMSD, maximum observed Cα displacements of 12 Å and 23 Å, and overall bend angles of 3.3° and 8.9° about the interface (**Fig. 3e-f** and **Table S6**).

These results indicate that filaments composed of the liganded abortive complexes are more variable and hence more dynamic about the hexamer-hexamer interfaces in comparison to apo filaments. Ligand binding triggers subtle conformational changes that alter filament assembly interfaces, which ultimately influence filament dynamics and stability.

### The hexamer-hexamer contact restrains the full range of conformational changes that accompany formation of the closed state abortive complex

Upon binding substrates, bGDH subunits undergo a conformational change that shifts the position of the NAD binding domain towards the Glu binding domain. This shift closes the active site around bound ligands, with accompanying conformational changes in the antenna region (including the long antenna helix and nearby residues, **Fig. 4a**). However, these structural changes are dampened in the liganded di-hexamer (green, **Fig. 4a**). To quantify the changes, we plot the per-residue RMSDs between apo vs. liganded states for the mono-(teal) and for the di-hexamers (salmon) in **Figure 4b**. The difference between these two lines (ΔRMSD) is shown in black. If the conformational transition upon ligand binding is substantial, we would accordingly observe large RMSDs. Conversely, if the conformational transition is small, we would observe lower RMSDs. The plot shows that the degree of conformational change resulting from ligand binding in filamentous di-hexamer assemblies is dampened compared to that of the mono-hexamers by an amount indicated by the black line.

**Figure 4.**
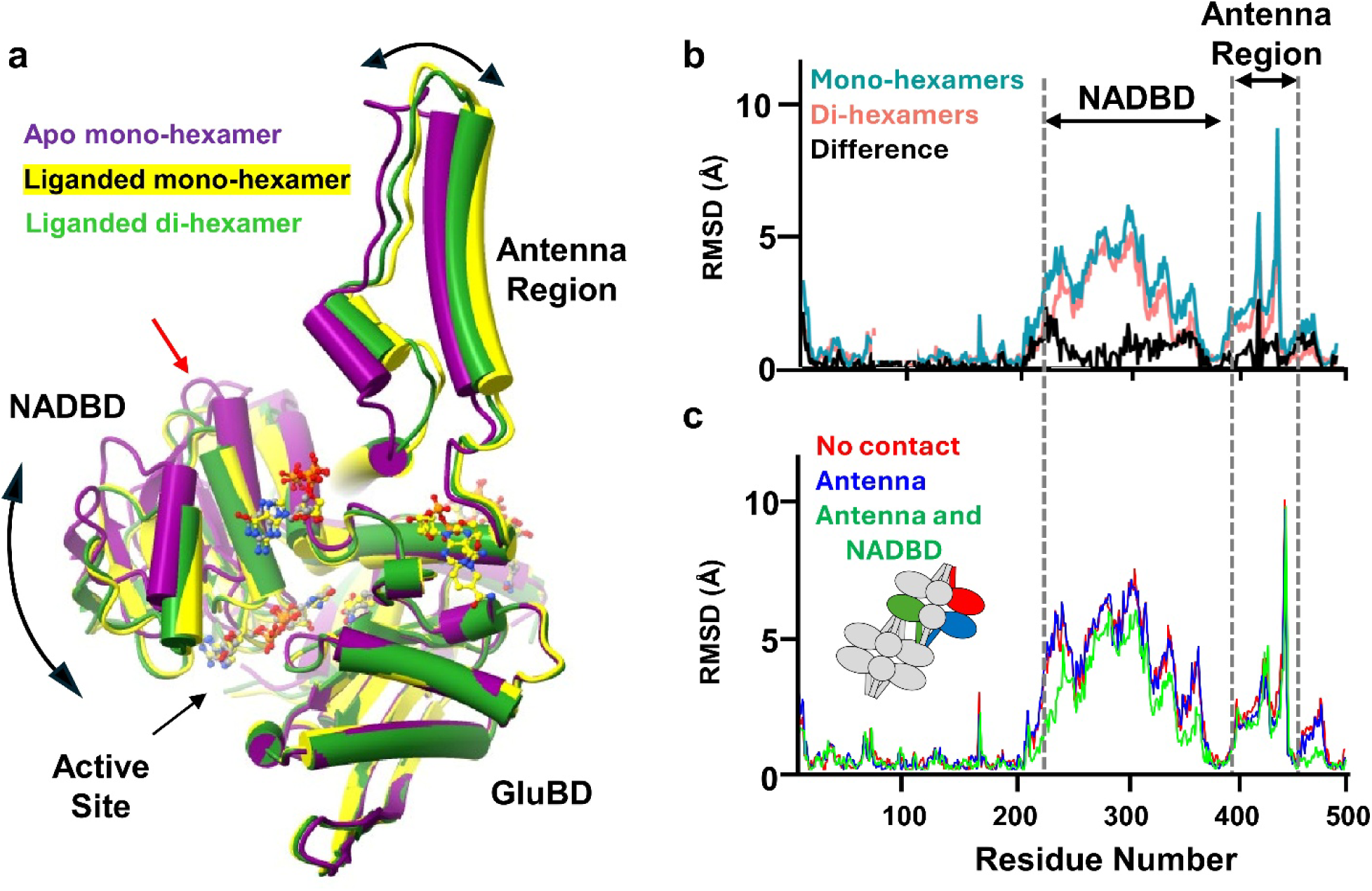
Hexamer-hexamer contacts dampen conformational changes associated with ligand binding. **a,** Comparison of a single subunit of bGDH from the apo mono-hexamer (dark purple), liganded mono-hexamer (yellow), and liganded di-hexamer (green). The red arrow shows the location of the contact site with the neighboring hexamer in the di-hexamer structures. NADBD, NAD binding domain, GuBD, Glu binding domain. The curved arrow shows the shift in the NAD binding domain beween apo and liganded structures. Superposition performed with Cα of the GluBD. **b,** RMSD (Å) between apo and liganded states in mono-(teal) and di-hexamer (salmon) structures in a single subunit using Cα of the GluBD (residues 5-208, 474-496, 369-390) in the superposition. The difference between these two lines (black) shows that di-hexamers display smaller RMSDs because hexamer-hexamer contacts hinder conformational changes associated with ligand binding. A subunit engaged via both its NAD binding domain and antenna in the hexamer-hexamer contact was used in the superposition. **c,** As in (b), but using apo and liganded di-hexamer structures, and the GluBD of all six hexameric subunits in the superposition. Red indicates RMSD for subunits with no hexamer-hexamer contacts, blue for antenna-only contacts, and green for both NADBD and antenna contacts. The lower values for the latter subunit indicate more restricted conformational changes upon ligand binding. **(Inset)** Cartoon illustrating the position of subunits for which RMSDs are plotted.

Dampened conformational changes between apo and liganded states for filamentous bGDH assemblies are not symmetric throughout hexamers and depend on the number of contacts at the hexamer-hexamer interface. The lowest RMSD values (corresponding to the largest dampening) are observed in the subunit of one hexamer that makes two contacts to the neighboring hexamer (green, **Fig. 4c**). Intermediate dampening is observed in subunits that engage via only their antenna helix (blue, **Fig. 4c**). The highest RMSD values (and therefore the least dampening of conformational changes) are observed in subunits that do not participate in hexamer-hexamer contacts (red, **Fig. 4c**). Therefore, contacts between hexamers that stabilize bGDH filaments restrict the conformational changes associated with ligand binding due to the restriction on the shifting of the NAD binding domain.

## DISCUSSION

Here, we set out to understand the role of filamentation in mammalian GDH function. We confirmed that bGDH forms linear, filamentous self-assemblies at concentrations that are typical within mitochondria^3^. Four high-resolution cryo-EM structures demonstrate the basic mechanism through which bGDH hexamers self-associate via two reciprocal contact points each involving an NAD-binding domain of one hexamer and the antenna tip of the other. This self-association is the basis for filament formation and breaks the inherent 3-fold symmetry of individual hexamers (**Fig. 1b-e, 2d-f,** and **Fig. S13**). Binding of ligands that result in the abortive complex shift bGDH into a closed, inhibited conformation that weakens the hexamer–hexamer interactions, yielding more flexible and shorter filaments. The persistence of the hexamer-hexamer contact in crosslinked di-hexamer complexes results in a dampened structural response to ligand binding in subunits most closely involved in the contact interfaces.

### Filamentation opposes the formation of the inhibited abortive complex

To fully close the active site around bound ligands, the NAD binding domain must shift towards the Glu binding domain, a transition that also alters the hexamer-hexamer interface. The full range of conformational changes associated with ligand binding, particularly within the antenna region and parts of the NAD binding domain, is resisted in the presence of structural constraints from filamentation. This is clearly evidenced through RMSD analyses between the individual subunits of apo and liganded complexes (**Fig. 4** and **Fig. S11-13**). Subunits that directly engage in hexamer-hexamer contacts exhibit the most dampened conformational changes associated with active site closing, whereas subunits that do not engage in hexamer-hexamer contacts show conformational changes associated with ligand binding that are similar to those found in the mono-hexamer structure.

GTP binding stabilizes the closed and inactive state of bGDH, preventing release of abortive complex ligands as well as access of new substrates to the active site^28,29,42–44^. GTP binding also results in a shift in the equilibrium towards hexameric and away from filamented bGDH^17,24,33^. Interestingly, kinetic studies show faster rate constants for hexamer dissociation from filaments in the presence of GTP than in its absence^33^, and it is known that as many as 2 moles of GTP can bind per mole hexamer without inducing dissociation of bGDH from filaments^45^. These observations suggest that GTP binds hexamers in filamentous bGDH assemblies, and only after more than two of the six GTP binding sites are occupied do hexamer-hexamer contacts weaken sufficiently for filaments to dissociate. Our structural studies show that saturation of all GTP binding sites (in conjunction with NADH and Glu binding) within hexamers weakens hexamer-hexamer interactions. The bGDH complexes used in our studies included all ligands prior to the introduction of the crosslinking agent, implying that we have captured either hexamers in a weakened state of interaction or hexamers that are progressing to full dissociation from filamentous assemblies. In either case, the binding of these abortive complex/GTP ligands has altered and weakened the interactions between bGDH hexamers within filaments.

These collective observations indicate that filamentation through hexamer self-association constrains the conformational changes associated with formation of the inhibited, liganded complex, particularly in those subunits at the hexamer-hexamer interface. This would be expected to weaken GTP binding, consistent with experimental observations^24^. Conversely, we find that the binding of GTP, in the presence of Glu and NADH, weakens the interactions that maintain the filamentous state, which is expected to shift the equilibrium toward filament dissociation into hexamers, also consistent with observations^17,24,33^. Together, these results show that filamentation and GTP binding are mutually antagonistic: filament assembly limits the formation of the GTP-inhibited state, while GTP binding in the liganded complex promotes filament destabilization and favors the hexameric enzyme.

### Model for the role of filamentation in GDH function

bGDH forms a hexameric oligomer with D3 symmetry (**Fig. 5a**). Each of the six bGDH subunits contains an active site located between an NAD binding domain and a glutamate binding domain, as well as one antenna helix that assembles into a trio of helices forming an antenna on either side of the hexameric complex. Our biochemical and structural data show that bGDH hexamers assemble into linear filaments via reciprocal contacts formed by the antenna of one hexamer and an NAD binding domain of another hexamer (**Fig. 5b**). Formation of the GTP stabilized, abortive complex requires a shift in the position of the NAD binding domain (red arrow, **Fig. 5b, Movie S7**), which disrupts the hexamer-hexamer contacts (black arrows, **Fig. 5b**). However, formation of such an abortive complex is resisted by the hexamer-hexamer contacts found in bGDH filaments. In agreement with published kinetic analyses^24,31^, we propose a working model for the role of filamentation in regulating bGDH activity. When cells reside in a low energy state, such as when the ratio of ADP/GTP is high, bGDH forms filaments in the mitochondria, while ADP allosterically activates bGDH to catalyze oxidative deamination of glutamate and generate αKG, with NAD(P)H being a by-product of the reaction. While αKG is metabolized to fuel the Krebs cycle, the reduced coenzymes can be used for further energy production (NADH) or biosynthesis (NADPH). This reaction is sustained until cells return to a high energy state characterized by a low ADP/GTP ratio, whereby the activity of bGDH is reduced by the inhibitory activity of GTP. However, filamentation results in a level of resistance to the complete inhibition of bGDH, since disruption of hexamer-hexamer interactions would become necessary. Thus, filamentation could allow bGDH to remain catalytically competent (albeit at a reduced level) at concentrations of GTP which would otherwise fully inhibit non-filamentous GDH. This may be advantageous in cells that must continue to oxidize glutamate and generate αKG/NADPH despite a high-energy state, for example, to support biosynthesis or glutamate clearance.

**Figure 5.**
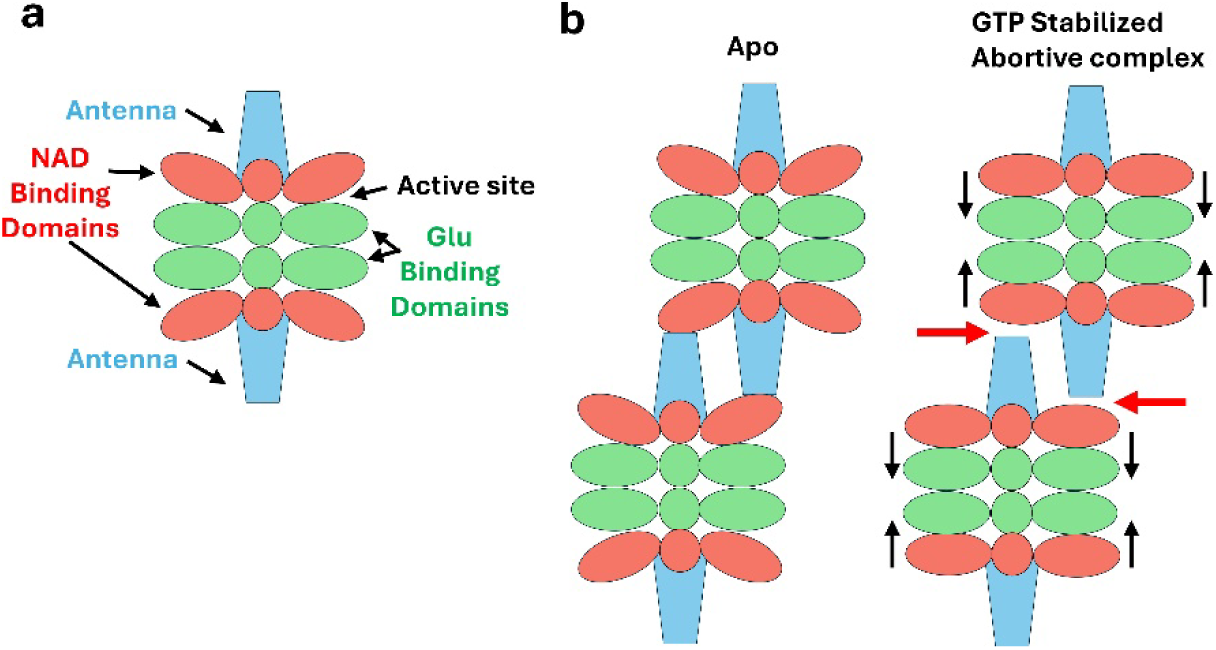
Model for how hexamer-hexamer contacts within filaments hinder the formation of the abortive complex. **a,** Schematic of mono-hexameric bGDH. Each hexamer contains two antenna at either end, six NAD binding domains (NADB), six glutamate binding domains, and six active sites. **b**, The interaction between bGDH hexamers found in filamentous assemblies. The antenna of one hexamer contacts a single NADBD of another hexamer. Formation of the abortive complex requires shifting of the NADBD to close the active site around the bound ligands glutamate and NADH (black arrows). This conformational change disrupts hexamer-hexamer contacts (red arrows). Hence, the formation of the abortive complex is resisted by the loss of contact energy in the hexamer-hexamer contacts, requiring higher concentrations of GTP to achieve the same inhibition observed in mono-hexameric bGDH.

This concept is exemplified by the primate neural isoform GDH2 that has evolved a near-complete insensitivity to GTP, allowing glutamate oxidation even when GTP levels are sufficient to inhibit the housekeeping enzyme GDH1^2,46^. By analogy, we propose that filamentation of bGDH provides a structural mechanism to dampen GTP/NADH inhibition, enabling bGDH to continue supplying αKG and NADPH in contexts where glutamate oxidation and biosynthesis or glutamate clearance are still required despite a high-energy state. Control of bGDH filamentation could in turn fine tune GTP inhibition of GDH activity to meet specific cellular demands.

The strongest effect on bGDH conformation that would be expected to influence abortive complex formation and GTP binding is found in the subunit engaged with hexamer-hexamer contacts via its NAD binding domain. How this would affect the other five subunits of the bGDH hexamer depends on the level of cooperativity between subunits within the hexamer. Evidence from activity assays indicate that there is inter-subunit cooperativity with regard to coenzyme binding^47–49^, and structural studies show some degree of coordination between subunits connected by the formation of the antenna structure found at either end of the hexamer^30,50^. Cooperativity in conformational transitions between individual subunits would extend any dampening effects mediated by hexamer-hexamer interactions beyond the immediate NAD binding domain and to other subunits within the hexamer.

Finally, we determined the structure of bGDH in the inhibited, abortive complex state which exhibits a closed active site, but closure of the active site is expected to also occur during the normal bGDH reaction cycle during the conversion of substrates to products. Published studies indicate that there is no effect of bGDH filamentation on its basal catalytic rate in the absence of allosteric effectors^51,52^. Structures of human or bGDH with matched substrates or products have yet to be reported, and therefore it is not known if the conformational state of such a complex would differ significantly with that of the abortive complex. If similar, disruption of hexamer-hexamer interactions during the normal catalytic cycle would also be expected but may be transient, with compensating effects on rates of active site closing and opening resulting in little to no effect on the overall reaction rate. Alternatively, the closed conformation during a normal catalytic cycle may differ from that of the abortive complex in such a way to allow full hexamer-hexamer contacts to remain intact. Future studies will be necessary to investigate hexamer-hexamer interactions during the normal reaction cycle, as well as to address the potential for cooperativity between subunits in the context of bGDH filamentation.

### Limitations of the study

To stabilize di-hexamers of bGDH for structural studies, we used chemical crosslinking in the presence of high sodium sulfate concentrations. We cannot control the precise state in which the structures are captured, since filamentation appears to be a dynamic process, even more so when ligands weaken the hexamer-hexamer interface. Indeed, we may be averaging a multitude of intermediate states, captured through chemical crosslinking. However, the resulting assemblies from chemical crosslinking match those seen under physiological conditions (**Fig. S2**). Moreover, it is clear that the overall structural transitions are significant, leading to large-scale changes at the level of protein backbones and spanning entire interfaces. Thus, we believe that we are capturing bona fide on-pathway intermediates that broadly serve to illuminate a complex mechanism of filament-mediated regulation of enzyme activity.

## ONLINE METHODS

### Sedimentation equilibrium Analysis (AUC)

Sedimentation equilibrium experiments were conducted using a Beckman Coulter Optima multi-wavelength analytical ultracentrifuge equipped with a Ti-50 rotor. Bovine GDH (bGDH, GLUD1, EC 1.4.1.3) was obtained from Sigma-Aldrich (cat. no. G2626), stored at −20°C, and used without further purification. Three concentrations (1.5, 1.0, and 0.5 mg/ml) of bGDH were prepared from dilution into buffer A (50 mM Tris-HCl, 150 mM NaCl, pH 7.5 at room temperature) followed by extensive dialysis against the same buffer. These samples were loaded into three chambers of a six-chamber centerpiece, with the remaining three chambers filled with buffer to serve as a reference. The cells in the rotor were allowed to equilibrate at 25°C for 1 hour before starting the centrifugation. Centrifugation speeds were increased by 2,000 rpm increments, starting from 2,000 rpm and progressing through 4,000, 6,000, and 8,000 rpm. Equilibrium was confirmed when the RMSD between consecutive scans (taken every 2 hours) reached 0.07 or lower (typically after 72-96 hours). The equilibrium data from the 4,000, 6,000 and 8,000 rpm scans were analyzed globally using SEDPHAT^53,54^ and a multiple species model with one species fit at the molecular weight of the hexamer, and two other species with molecular weights allowed to be varied to best fit the data. The partial specific volume of bGDH was calculated from its amino acid sequence using SEDNTERP^55^ giving a value of 0.735 mL/g. Buffer density of and viscosity were also calculated using SENTERP giving values of 1.0009 g/ml and 0.01240 Poise, respectively.

### Determination of bGDH molecular weight distribution by mass photometry

The molecular weight distribution of bGDH (at 1 and 5 μg/ml) was performed using a OneMP mass photometer (Refeyn, Inc.). Bovine serum albumin (BSA; monomer: 66 kDa, dimer: 132 kDa) (Sigma Aldrich™ cat. no. 05470) and GroEL (801 kDa, kindly provided by Prof. Michael Marty of the University of Texas, Austin) were used as molecular weight standards to generate a calibration curve. bGDH was diluted in buffer A before measurement. Mass Photometry data was recorded over one minute and analyzed using DiscoverMP software (Refeyn, Inc.). Measurements were performed in triplicate. The percentage of HMWS (high molecular weight species, i.e. species with 2 or more bGDH hexamers) was based on the mass of species, meaning that the total mass of bGDH found in particles with 2 or more hexamers divided by the total mass including that of individual hexamers.

### Determination of bGDH molecular weight distribution and the effect of buffer conditions by negative stain electron microscopy

Sample preparation proceeded as follows: for the low ionic strength buffer (**Fig. S2a**), bGDH (50 µg/mL) was incubated in 25 mM Tris-HCl, 50 mM NaCl, pH 7.0 at 4 °C for 1 hour prior to imaging. The sample crosslinked in low ionic strength buffer (**Fig. S2b**) was prepared by incubating bGDH (50 µg/mL) in 20 mM sodium phosphate, pH 7.0 and 0.05% glutaraldehyde (Sigma-Aldrich cat. no G5882) at room temperature for 10 minutes before quenching with 50 mM Tris-HCl pH 7.5 prior to imaging. The sample in high sodium sulfate concentration buffer (**Fig. S2c**) was prepared with bGDH at 1 mg/ml incubated at room temperature in 20 mM sodium phosphate, 1 M sodium sulfate, pH 7.0 for 1 hour followed by dilution to 50 μg/ml before imaging. Finally, bGDH crosslinked in the high sodium sulfate buffer (**Fig. S2c**) utilized bGDH at 1 mg/ml in 20 mM sodium phosphate, 1 M sodium sulfate, pH 7.0 mixed with 0.01% glutaraldehyde at room temperature for 1 hour followed by quenching with 50 mM Tris-HCl pH 7.5 and dilution to 50 μg/ml bGDH before imaging.

For imaging of samples, carbon lined copper grids (CF300 Cu grids, VWR cat. no. 103303-538) were glow-discharged using PELCO glow discharge at 15 mA current for 30 seconds. Samples were incubated on grids for 1 minute, then washed three times with water followed by incubation in 1-4.5% (w/v) uranyl formate for 1 minute. The uranyl formate was subsequently blotted using Whatman blot paper, and the grids were air dried at room temperature for 5 minutes. Grids were imaged using a 120 kV FEI Tecnai G2 Spirit BT transmission electron microscope with AMTXR80 (8MP) digital CCD camera at the University of Arizona. Oligomers of bGDH were identified in EM images as those showing very close proximity of bGDH hexamers in a roughly linear arrangement. Particles were counted and binned according to the number of hexamers per particle.

### Negative stain imaging of apo and liganded bGDH samples (Fig. S14)

Negative staining and imaging were performed on apo and liganded samples as follows: for apo samples, 0.5 mg/ml bGDH was incubated in a buffer containing 25 mM Tris-HCl, pH 8.0, 50 mM NaCl, 0.75 M ammonium sulfate. For liganded samples, the treatment was identical but with the addition of 5 mM glutamate, 0.4 mM NADH, and 0.9 mM GTP. Samples were applied to carbon-coated grids (Electron Microscopy Sciences) which had been glow-discharged using a PELCO glow discharger at 15 mA current for 30 seconds, and incubated for 1 minute, then washed three times with water, and finally incubated with 1% (w/v) uranyl formate for 1 minute. The uranyl formate was then blotted using Whatman blot paper, and the grids air dried at room temperature for 5 minutes. Grids were imaged using a 200 kV FEI Talos microscope with a 4k x 4k Ceta CCD camera.

### Sample preparation for Cryo-EM

bGDH filaments without ligands were assembled by incubating 1 mg/mL bGDH with a buffer containing 1 M sodium sulfate, 20 mM sodium phosphate buffer, pH 7.0 for one hour at room temperature. Crosslinking was then performed by adding glutaraldehyde (Sigma-Aldrich cat. no. G5882, prepared in 20 mM sodium phosphate pH 7.0 buffer) to a final concentration of 0.01% in the sample and incubating for one hour at room temperature before quenching with 50 mM Tris-HCl pH 7.5. The sample was then centrifuged for 2 min at 10,000xg and the supernatant loaded onto a Superdex 200 10/300 GL (GE) size-exclusion column that had been pre-equilibrated with a buffer containing 0.2 M sodium sulfate, 50 mM sodium phosphate, pH 7.5@4°C. Fractions (1 ml) were collected, concentrated, checked by negative stain EM, and those fractions containing filaments used for cryo-EM grid preparation.

For ligand-bound bGDH filaments, 1 mg/mL bGDH was mixed with 0.1 mM NADH, 0.1 mM GTP, and 10 mM glutamate in a buffer containing 20 mM sodium phosphate, 150 mM NaCl, and 600 mM sodium sulfate, pH 7.5 for 15 minutes at 30°C. The crosslinker dimethyl pimelimidate (DMP)(Thermo Scientific™ cat. no. 21666) was then added to a final concentration of 50 mM in 3 increments 15 minutes apart. The sample was then further incubated at 30°C for one hour before quenching with 50 mM Tris-HCl pH 7.5. Large aggregates were removed by centrifugation for 2 min at 10,000xg, and the supernatant loaded onto a Superdex 200 10/300 GL (GE) size-exclusion column that had been pre-equilibrated with buffer containing 15 mM sodium phosphate, 100 mM NaCl, 200 mM sodium sulfate and 5 mM glutamate at pH 7. Appropriate peak fractions were collected and concentrated to 1 mg/ml, then centrifuged at 10,000xg for 1 minute to remove large aggregates and imaged by negative stain EM. Fractions containing filaments were mixed with additional NADH and GTP to final concentrations of 0.4 mM NADH, 0.9 mM GTP and used directly for cryo-EM grid preparation. For both apo bGDH and ligand-bound bGDH samples, 2.5 μl of 0.5 mg/mL GDH samples were applied to plasma cleaned R1.2/1.3 gold UltrAuFoil grids, Au 300 mesh (Quantifoil), then frozen using a manual plunger in the cold room at 4 °C and 90% humidity. These grids were clipped and then subsequently stored in liquid nitrogen until data acquisition.

### Cryo-electron microscopy data collection

Both the apo bGDH and the ligand-bound bGDH datasets were collected using a Titan Krios transmission electron microscope (Thermo Fisher Scientific) operating at 300 KeV, equipped with a K3 quantum director (Gatan) with a GIF (Gatan Imaging Filter) BioQuantum energy filter with a slit width of 15 eV. The movies were recorded at a magnification of 105kx, corresponding to a pixel size of 0.822 Å/pixel in nanoprobe EF-TEM mode. Data collection was performed in an automated manner using the EPU software version 2.12 from Thermo Fisher Scientific. The total fluence was 40.4 e^-^/Å^2^ at a rate of 9.05 e^-^/pix/s for apo bGDH dataset, while the total fluence was 44.8 e^-^/Å^2^ at a rate of 10.1 e^-^/pix/s for ligand-bound bGDH dataset. To address issues with preferred orientation and avoid anisotropic resolution, the stage was tilted to 40° and 50° during automated data collection for both datasets^56–58^.

### Cryo-electron microscopy data processing

The same workflow for data processing using cryoSPARC^38^ and Relion^59^ was applied to both apo and ligand-bound bGDH datasets, unless otherwise noted. Movies were imported into cryoSPARC for motion correction and patch-based contrast transfer function (CTF) estimation. For the apo bGDH dataset, a 2D template was generated after an initial round of particle selection using the Blob Picker tool and 2D classification, both implemented in cryoSPARC. Subsequently, the 2D template was used for template-based particle selection. The following procedures were used for reconstruction of a hexamer map: the particles were extracted with a box size of 320, and Fourier cropped to 256 pixels after inspection of particle picking. Reference-free 2D classification was used to clean the particle stack, and the best 2D class averages were selected based on the appearance of particles containing recognizeable protein features. After multiple iterations of 2D classification, the best classes were selected to generate 3 *ab initio* reconstructions. The best *ab initio* reconstruction that contained features consistent with the known bGDH structure was then subjected to homogeneous refinement. We then performed multiple rounds of iterative heterogeneous refinement using more than three classes, followed by homogeneous refinement of the particles contributing to the highest-resolution reconstructions. D3 symmetry was applied throughout these procedures. The best volume and its corresponding particles from the last heterogeneous refinement were selected to perform a homogeneous refinement with D3 symmetry to improve the map. Subsequently, the particles were imported into Relion for particle polishing^59^ using default parameters. The polished particles were then imported back to cryoSPARC to perform per-particle CTF refinement, followed by a homogeneous refinement with D3 symmetry. Iterative Bayesian polishing in Relion and homogeneous refinement with D3 symmetry in cryoSPARC was repeated until no further improvements in map resolution were observed, as assessed using the Fourier shell correlation (FSC). The following procedures were used for reconstruction of the di-hexamer map. Particles were re-extracted with a box size of 512, then Fourier cropped to 320 pixels after inspection of particle picking. The same data processing workflow was performed for the di-hexamer map, with the exception that all refinement operations were performed with C1 symmetry. The poor density of the second hexamer in the di-hexamer reconstruction was better resolved by performing particle subtraction and masked local refinement.

For the ligand-bound bGDH dataset, only two major differences in the data processing workflow were used. First, to reconstruct the di-hexamer map for ligand-bound bGDH, the particles were re-extracted with a box size of 512, and Fourier cropped to 384 pixels after inspection of particle picking. Second, instead of performing Bayesian polishing in Relion, the movement trajectories of each particle were calculated using the Reference Based Motion Correction program^60^ which is implemented in cryoSPARC and is the cryoSPARC equivalent to the Relion Bayesian polishing approach. The processing workflows for both datasets are summarized in **Figures S4** and **S8**. The local resolution was calculated in cryoSPARC. The 3D FSC^56^ was obtained using the 3D FSC server and the sampling compensation function^61,62^ was calculated in cryoSPARC.

### Cryo-electron microscopy structure building and refinement

We used an atomic model of bGDH derived from an AlphaFold3 prediction and the full, unprocessed sequence of bGDH to build and refine the apo bGDH hexamer model, and hence residue numbering in the coordinate files corresponds to the initially translated and unprocessed protein. The apo bGDH hexamer model was fit into the apo bGDH map using Chimera^63,64^, and minor adjustments were made using Coot^65^. Subsequently, we performed one round of real-space refinement within the Phenix (https://phenix-online.org/) suite^66,67^. The models were then iteratively adjusted in Coot and refined in Phenix, and the statistics were examined using Molprobity^68,69^ until no further improvements were observed. The apo bGDH di-hexamer model was built using the same procedure, with the addition of the second bGDH hexamer. The ligand-bound hexamer and di-hexamer models were built using the respective apo bGDH models as references, and the same refinement and adjustment procedures as used for the apo models. The ligands were added using tools in Coot^70^, and the water molecules were built using the Douse program in Phenix for high-resolution maps. Water molecule occupancy was inspected in Coot manually using the validation features. The final models were also evaluated using map/model FSC analysis and using EMRinger^71^ to compare the fit of the model backbone into the cryo-EM map. The model statistics showed good geometry and matched the cryo-EM reconstruction. Refined coordinates (9ZQR, apo di-hexamer, 9ZQS, apo mono-hexamer, 9ZQT, liganded di-hexamer, 9ZQU, liganded mono-hexamer) and maps (EMD-74576, composite map of apo di-hexamer, EMD-74574, consensus map of apo di-hexamer, EMD-74575, constituent map A of the apo di-hexamer, EMD-74577, apo mono-hexamer map, EMD-74581, composite map of liganded di-hexamer, EMD-74578, consensus map of liganded di-hexamer, EMD-74579, constituent map A of the liganded di-hexamer, EMD-74580, constituent map B of the liganded di-hexamer, EMD-74582, apo mono-hexamer map) have been deposited in the RCSB and EMDB databases, respectively.

### 3-D Flexibility Analysis

We generated new dimer of hexamers particle stacks for 3D flexibility analysis using 3DFlex^41^ implemented in CryoSPARC (v4.7.0, Structura Biotechnology) using the cryo-EM data described above. First, we optimized particle picking using a combination of topaz^72^ (v0.2.5a) and template picking in cryoSPARC (v4.7.0). Second, we cleaned the initial particle stacks using 2D classification, 3D classification with alignment, and focused 3D classification. Importantly, we Fourier cropped particles (from 540 pixel to 128 pixel box sizes) and limited the alignment resolution to 25 Å to prevent overfitting to a single subunit of the dimer and to encourage a diversity of conformations in the resulting particle stacks. The resulting electron density maps were refined to the Nyquist limit at 7.22 Å resolution. We used the coulombic potential maps and particle stacks to train a 3DFlex model. Last, we interpreted the learned 3D flexibility of by generating deformations of the reference map using the 3DFlex model. We manually inspected the resulting maps to generate movies capturing motions of apo and liganded bGDH. To calculate bend angles captured in the conformational ensemble, we relaxed atomic models derived from high resolution structural analysis into two separate low resolution maps using Isolde (v1.9)^73^ implemented in ChimeraX (v1.9, UCSF). The two maps represented the two extremes of conformational motions learned by 3DFlex. We modified a script called modevectors (by Sean M. Law, University of Michigan) to calculate and plot Cα displacement in the resulting atomic models in PyMOL (v3.1.6, Schrodinger). We calculated RMSD between the two models after global alignment using PyMOL (v3.1.6). We calculated dihedral bend angles of the resulting models using Chimera (v1.17.3, UCSF).

### RMSD Calculations and Movies

Root mean square displacements (RMSD) between Cα in comparisons between structures were calculated using default settings in the MatchMaker^74^ application in Chimera^75^. Several cycles of pruning were performed to achieve the best alignment. RMSD was then calculated for all Cα pairs present in both structures used in the comparison. **Movies S1-S2** were prepared in 3DFlex software, and **Movies S3-S7** were made with Pymol software v. 1.8.0.3 (Schrodinger, Inc.), the morphs were prepared in ChimeraX v1.8 (UCSF Software), and video editing performed with OpenShot v3.4.0.

## ACKNOWLEDGEMENTS

This work was supported by the National Science Foundation under Grants MCB-1934291 (N.C.H. and D.L.); DBI-2018942 (N.C.H.), and MCB-2048095 (D.L), the National Institution of Health under grants T32GM136536 (N.I.D.), F32GM148049 (T.S.S.), and F31AI189273 (A.R.G.), the NOMIS Foundation fellowship (T.S.S.), Hearst Foundations Developmental Chair (D.L.), and the University of Arizona Research, Innovation & Impact (RII) and Technology Research Initiative Fund/Improving Health and Access and Workforce Development (N.C.H.). Some of the transmission electron microscopy imaging was performed at the University of Arizona, ORP Imaging Cores - Electron facility, RRID:SCR_023279 with assistance from Dr. Paula Tonino. The FEI Tecnai G2 Spirit BT TEM was supported by the grant from the NIH S10-OD011981.

