## Supplementary Tables and Figures for "Structural Mechanism of Filamentation Induced Dampening of GTP Inhibition of Glutamate Dehydrogenase"

**Supplementary Information for**
### Structural Mechanism of Filamentation Induced Dampening of GTP Inhibition of Glutamate Dehydrogenase

**Zelin Shan[1†], Noura I. Darwish[2†], Andres Rivero-Gamez[1,3], Timothy S. Strutzenberg[1], Dmitry Lyumkis[1,3*], Nancy C. Horton[2*]**

[†]These authors contributed equally to this work.
[1] The Salk Institute of Biological Sciences, La Jolla, CA, USA 92037
[2] Department of Molecular and Cellular Biology, University of Arizona, Tucson, AZ, USA 85721
[3] Department of Molecular Biology, School of Biological Sciences, University of California, San Diego, La Jolla, CA, USA 92093

[*]To whom correspondence should be addressed:

#### Supplementary Tables

**Table S1. Molecular weight analysis of bGDH by mass photometry**

| Species | Counts in Measurement 1 | Counts in Measurement 2 | Counts in Measurement 3 | Percentage of Total by Mass[1] |
|---|---|---|---|---|
| | | **1.25 μg/ml bGDH** | | |
| **Single Chain** | 256 | 111 | 231 | 5±2% |
| **Trimer** | 63 | 15 | 24 | 3±2% |
| **Hexamer** | 553 | 666 | 478 | 88±5% |
| **2 Hexamers** | 16 | 8 | 9 | 3±1% |
| | | **5 μg/ml bGDH** | | |
| **Trimer** | 180 | 169 | 155 | 1.9±0.3% |
| **Hexamer** | 4380 | 3197 | 3676 | 86±1% |
| **2 Hexamers** | 337 | 258 | 262 | 13±1% |
| | | **10 μg/ml bGDH** | | |
| **Hexamer** | 10330 | 7520 | 5992 | 86±2% |
| **2 Hexamers** | 775 | 730 | 457 | 7±1% |

[1]The average and standard deviation were calculated by mass and from three independent measurements.

**Table S2. Hexamer assembly statistics of 50 μg/ml bGDH by negative stain TEM**

| Species | Image 1 | Image 2 | Image 3 | Percentage of Total by Mass[1] |
|---|---|---|---|---|
| **Hexamer** | 1856 | 2368 | 2100 | 40±5% |
| **2 Hexamers** | 1472 | 1056 | 1504 | 51±9% |
| **3 Hexamers** | 64 | 224 | 96 | 7±5% |
| **4 Hexamers** | 17 | 15 | 13 | 1.1±0.2% |

[1]The average and standard deviation were calculated from three independent measurements.

**Table S3. Molecular weight distribution from sedimentation equilibrium analysis of bGDH at 0.5 mg/ml and three centrifugation speeds**

| Species | 4 krpm | 6 krpm | 8 krpm | Percentage of Total by Mass[3] |
|---|---|---|---|---|
| **1 Hexamer[1] (330 kDa)** | 48% | 49% | 36% | 41±6% |
| **3 Hexamers[2] (909 Da)** | 49% | 53% | 61% | 54±6% |
| **7 Hexamers[2] (11.9 MDa)** | 3% | 5.5% | 3.6% | 4±1% |

[1]The molecular weight of this species was fixed at the predicted molecular weight of a single hexamer.
[2]The molecular weight of this species was fit by SEDPHAT(1).
[3]The average and standard deviation were calculated from the three different speeds.

#### Table S4. Cryo-EM data collection statistics

| | **bGDH apo** | | | **bGDH bound to ligands** | | | |
|---|---|---|---|---|---|---|---|
| Microscope | Thermo Fisher Titan Krios | | | Thermo Fisher Titan Krios | | | |
| Voltage (KeV) | 300 | | | 300 | | | |
| Camera | Gatan K3 | | | Gatan K3 | | | |
| Magnification | 105,000 | | | 105,000 | | | |
| Pixel size at detector (Å per pixel ) | 0.822 | | | 0.822 | | | |
| Total electron fluence (e-/Å$^2$) | 40.4 | | | 44.8 | | | |
| Fluence rate (e-/Å$^2$/s) | 9.05 | | | 10.1 | | | |
| Stage tilt during data collection (°) | 40 and 50 | | | 40 | | | |
| Number of frames per movie | 50 | | | 50 | | | |
| Defocus range (µm) | 1.0-3.0 | | | 1.0-2.6 | | | |
| Automation software | EPU | | | EPU | | | |
| Energy filter slit width (eV) | 15 | | | 15 | | | |
| Total movies collected (no.) | 3950 | | | 5176 | | | |
| Micrographs used (no.) | 3480 | | | 5176 | | | |
| Total extracted particles (no.) | 330,194 | 330,194 | 1,666,463 | 1,117,578 | 1,117,578 | 1,117,578 | 1,974,264 |
| | Di-hexamer consensus map | Focus-refined hexamer map | Mono-hexamer map | Di-hexamer consensus map | Focus-refined hexamer map 1 | Focus-refined hexamer map 2 | Mono-hexamer map |
| | (map1) | (map2) | (map3) | (map0$_L$) | (map1$_L$) | (map2$_L$) | (map3$_L$) |
| | EMD-74574 | EMD-74575 | EMD-74577 | EMD-74578 | EMD-74579 | EMD-74580 | EMD-74582 |
| Particles in final reconstruction (no.) | 149,135 | 79,910 | 132,297 | 97,554 | 68,334 | 97,554 | 705,197 |
| Global resolution (FSC 0.143, Å) | 2.6 | 2.9 | 2.5 | 3.4 | 3.5 | 3.3 | 2.2 |
| Map sharpening B factor (Å$^2$) | -59.7 | -62.0 | -108.0 | -90.8 | -86.7 | -86.3 | -95.2 |
| SCF | 0.957 | 0.975 | 0.949 | 0.963 | 0.960 | 0.964 | 0.953 |

**Table S5. Model composition, refinement and validation**

| | Di-hexamer (apo) | Mono-hexamer (apo) | Di-hexamer (Ligand-bound) | Mono-hexamer (Ligand-bound) |
|---|---|---|---|---|
| | PDB: 9ZQR | PDB: 9ZQS | PDB: 9ZQT | PDB: 9ZQU |
| Model composition | | | | |
| Non-hydrogen atoms (no.) | 46,047 | 23,325 | 47,760 | 24,616 |
| Protein residues (no.) | 5885 | 2,952 | 5916 | 2,946 |
| Ligands (no.) | | | 42 | 24 |
| GTP (no.) | | | 12 | 6 |
| Glutamate (no.) | | | 6 | 6 |
| NADH (no.) | | | 24 | 12 |
| Water (no.) | | 243 | | 772 |
| Model refinement | | | | |
| Initial models used (PDB#) | | Alphafoldserver | | |
| Refinement packages | Coot, Phenix | Coot, Phenix | Coot, Phenix | Coot, Phenix |
| Model-Map scores | | | | |
| Cross-correlation coefficient | 0.81 | 0.82 | 0.84 | 0.90 |
| Model resolution (Å) | 3.1 | 2.8 | 3.6 | 2.4 |
| FSC threshold | 0.5 | 0.5 | 0.5 | 0.5 |
| Mean B factors (Å$^2$ ) | **Composite map values** | | **Composite map values** | |
| Protein residues (no.) | 46.94 | 56.50 | 79.35 | 34.07 |
| Ligands (no.) | | | 80.72 | 33.89 |
| Water(no.) | | 38.56 | | 34.71 |
| RMSD from ideal values | | | | |
| Bond lengths (Å) | 0.004 | 0.002 | 0.005 | 0.004 |
| Bond angles (°) | 0.466 | 0.389 | 0.487 | 0.492 |
| Validation | | | | |
| MolProbity score | 1.48 | 1.34 | 1.49 | 1.40 |
| CaBLAM outliers (%) | 0.82 | 0.99 | 0.63 | 0.21 |
| Clashscore | 8.92 | 6.22 | 9.10 | 7.25 |
| Rotamer outliers (%) | 0.02 | 0.73 | 0.40 | 0.61 |
| C-beta outliers (%) | 0 | 0 | 0 | 0 |
| Ramachandran plot | | | | |
| Favored (%) | 98.31 | 98.10 | 98.13 | 99.56 |
| Allowed (%) | 1.69 | 1.90 | 1.87 | 0.44 |
| Outliers (%) | 0 | 0 | 0 | 0 |

**Table S6: 3D Flexibility Analysis for apo and liganded bGDH filaments**

| **Atomic model** | **Aligned hexamer RMSD (Å)** | **mobile hexamer RMSD (Å)** | **Global RMSD (Å)** |
|---|---|---|---|
| **Liganded di-hexamer** | 1.197 | 2.031 | 3.087 |
| **Apo di-hexamer** | 0.783 | 0.907 | 1.818 |

**Morphs between bGDH structures, Movies S1-S4 and Movie S7:** Movies were made with Pymol software v. 1.8.0.3 (Schrodinger, Inc.), the morphs were prepared in ChimeraX v1.8 (UCSF Software), and video editing performed with OpenShot v3.4.0.

**Movie S1** shows a morph between mono- and di-hexamers of bGDH, both in the apo state, showing little effect of the hexamer-hexamer contact on the bGDH conformation.

**Movie S2** shows a morph between apo and liganded mono-hexamers of bGDH which illustrates the change in conformation of the bGDH hexamer and its subunits related to ligand binding.

**Movie S3** shows a morph between mono- and di-hexamers of bGDH, both in the liganded state, which shows: 1) changes in conformation of residues at the hexamer-hexamer interface, and 2) a difference in the conformation of subunits involved in the hexamer-hexamer contact in liganded di-hexamers compared to a liganded mono-hexamer. This difference is believed to be due to dampening of the shift in the NAD binding domain associated with ligand binding as a result of the hexamer-hexamer contact.

**Movie S4** shows a morph between apo and liganded di-hexamers which illustrates the change in the conformation of subunits as well as at the hexamer-hexamer interface in di-hexamer structures resulting from ligand binding.

**Movies S5-S6:**
3DFlex models were trained on dimer-of-hexamer particle stacks from apo and liganded bGDH datasets, and then these models have been used to compute the learned ranges of motion in corresponding low-resolution maps. The resulting maps show a striking increase in flexibility in the liganded filamentous bGDH (**Movie S6**) compared to that in the apo filamentous bGDH (**Movie S5**).

**Movie S7** shows a morph between the apo bGDH di-hexamer and liganded bGDH mono-hexamers (superimposed onto each hexamer of the di-hexamer) which shows how the NAD binding domain must rotate away from the hexamer-hexamer interface in order to close the active site around the ligands.

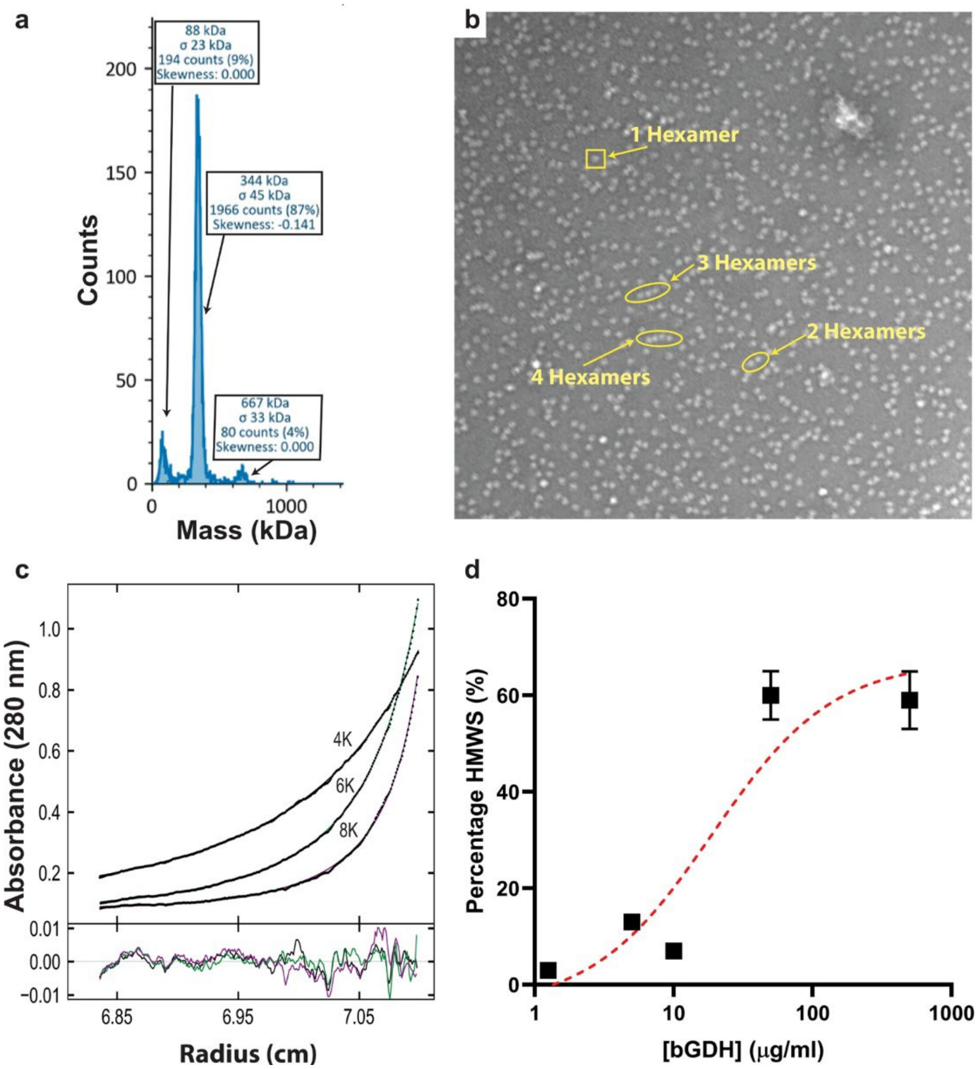

**Figure S1**. **bGDH forms filaments in a concentration dependent manner. a,** Mass photometry analysis of bGDH at 5 μg/mL, showing the distribution of oligomeric species, including hexamer (~344 kDa, theoretical MW=336 kDa), di-hexamer (~667 kDa, theoretical MW=672 kDa) and a peak corresponding to an unresolved mixture of monomeric and dimeric species (~88 kDa, theoretical MW=56 kDa for monomer, and 112 kDa for dimer). **b,** Representative negative stain EM image of 50 μg/mL bGDH incubated for 1 hour at 4°C in 25 mM Tris-HCl, 50 mM NaCl, pH 7.0. A single hexamer is outlined in a yellow box, and filaments of 2-4 hexamers are outlined in yellow ovals. **c,** Sedimentation equilibrium analysis of bGDH at 500 μg/ml. Fits to the data are shown as lines with their residuals shown in the lower plot. **d,** Summary of the mass photometry, EM, and sedimentation analysis plotted as the percentage (%) of the sample (by mass) which forms High Molecular Weight species (HMWS, filled squares, species with 2 or more hexamers) vs. the concentration of bGDH in μg/ml demonstrating a concentration-dependent oligomerization. The red dashed line is a fit to a binding isotherm giving a $K_{1/2}$ of 20 μg/ml.

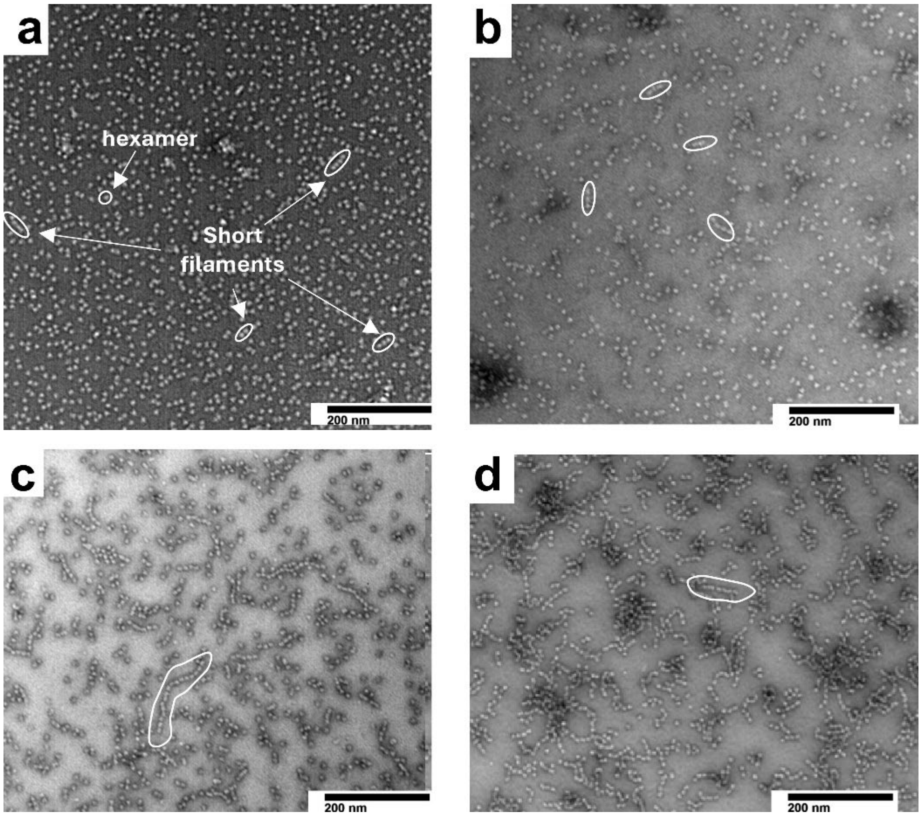

**Figure S2. Negative-stain TEM images of bGDH filaments with and without high ionic strength conditions and crosslinking**. **a,** bGDH (50 µg/mL) in low ionic strength buffer (25 mM Tris-HCl, 50 mM NaCl, pH 7.0) and incubated at 4 °C for 1 hour. Selected hexamer and filaments outlined in white. **b,** bGDH (50 µg/mL) in 20 mM sodium phosphate, pH 7.0, crosslinked with 0.05% glutaraldehyde (GA) at room temperature for 10 minutes. Selected filaments outlined in white. **c,** bGDH at 1 mg/ml in 20 mM sodium phosphate, 1 M sodium sulfate, pH 7.0 for 1 hour at room temperature followed by dilution to 50 µg/ml before imaging. One of the many long filaments is outlined in white. **d,** bGDH at 1 mg/ml in 20 mM sodium phosphate, 1 M sodium sulfate, pH 7.0 crosslinked with 0.01% GA at room temperature for 1 hour, then diluted to 50 ug/ml bGDH before imaging. One of the many long filaments is outlined in white.

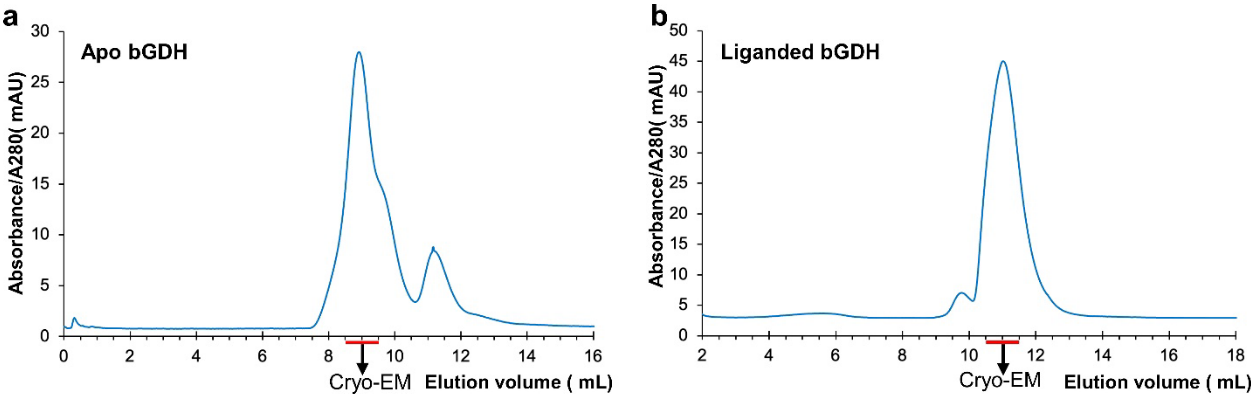

**Figure S3. Size exclusion chromatograms of bGDH samples used for cryo-EM. a,** Chromatogram of apo bGDH following treatment as described in Methods. The fractions used for cryo-EM are indicated. **b,** As in panel a, but for liganded bGDH.

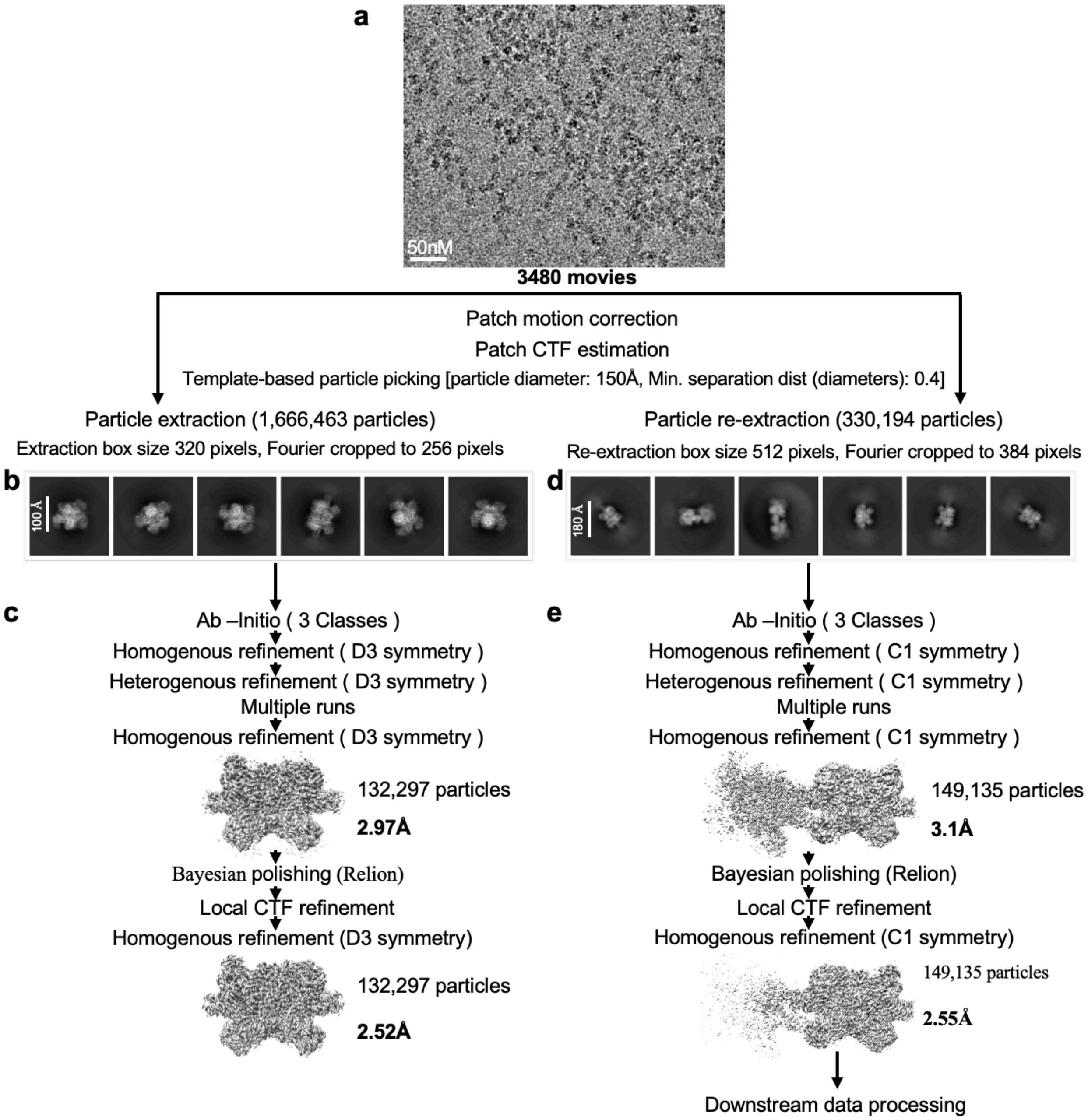

a
50nM
3480 movies
Patch motion correction
Patch CTF estimation
Template-based particle picking [particle diameter: 150Å, Min. separation dist (diameters): 0.4]
Particle extraction (1,666,463 particles)
Extraction box size 320 pixels, Fourier cropped to 256 pixels
Particle re-extraction (330,194 particles)
Re-extraction box size 512 pixels, Fourier cropped to 384 pixels
b
100 Å
d
180 Å
c
Ab –Initio ( 3 Classes )
Homogenous refinement ( D3 symmetry )
Heterogenous refinement ( D3 symmetry )
Multiple runs
Homogenous refinement ( D3 symmetry )
132,297 particles
2.97Å
Bayesian polishing (Relion)
Local CTF refinement
Homogenous refinement (D3 symmetry)
132,297 particles
2.52Å
e
Ab –Initio ( 3 Classes )
Homogenous refinement ( C1 symmetry )
Heterogenous refinement ( C1 symmetry )
Multiple runs
Homogenous refinement ( C1 symmetry )
149,135 particles
3.1Å
Bayesian polishing (Relion)
Local CTF refinement
Homogenous refinement (C1 symmetry)
149,135 particles
2.55Å
Downstream data processing

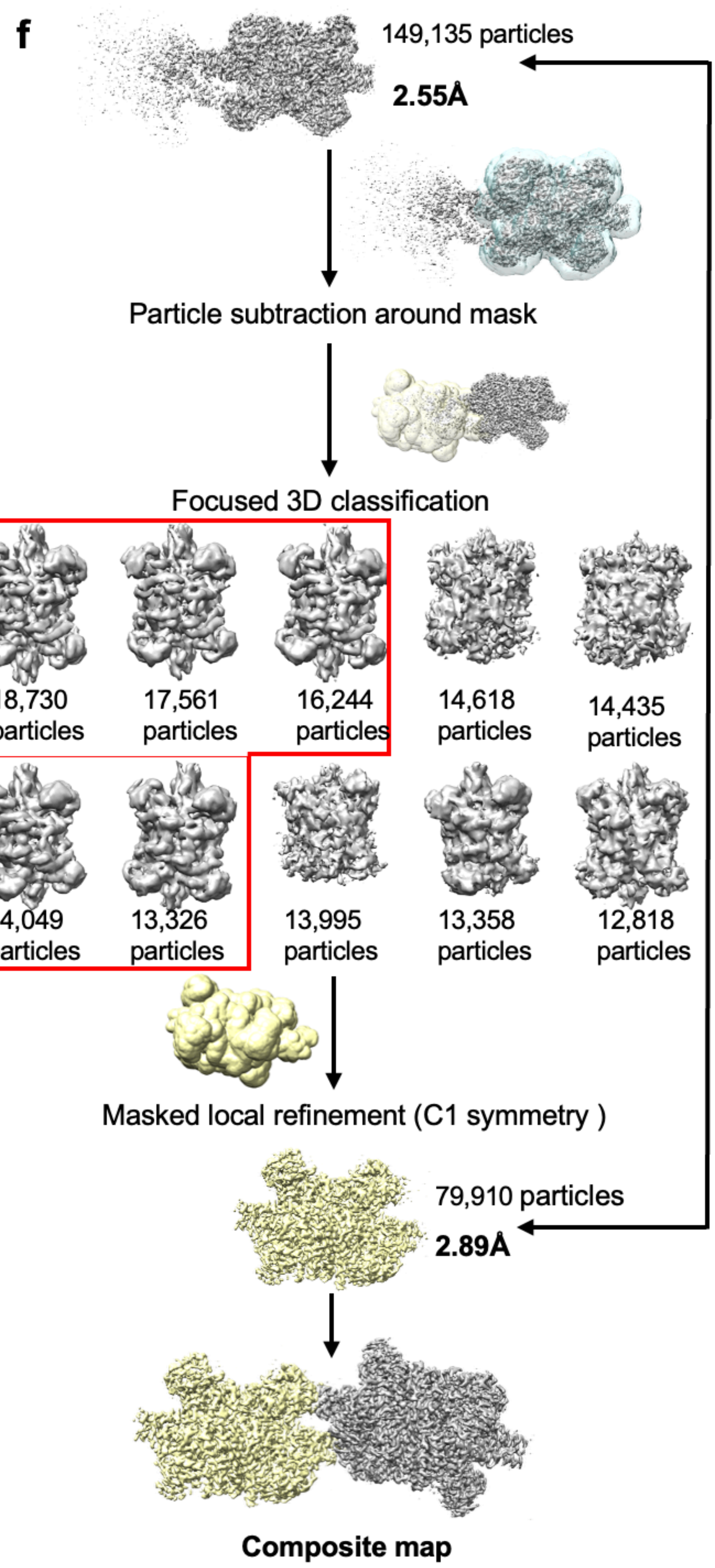
f
149,135 particles
2.55Å
Particle subtraction around mask
Focused 3D classification
18,730 particles
17,561 particles
16,244 particles
14,618 particles
14,435 particles
14,049 particles
13,326 particles
13,995 particles
13,358 particles
12,818 particles
Masked local refinement (C1 symmetry )
79,910 particles
2.89Å
Composite map

**Figure S4. Cryo-EM data collection and initial image processing of the apo bGDH dataset.** **a,** Representative denoised cryo-EM micrograph, scale bar is indicated. **b,** Representative 2D class averages from reference-free 2D classification showing the mono-hexameric particles, scale bar is indicated. **c,** Representative flowchart of the initial stage of imaging processing, ranges from 3D reconstruction to map refinement for mono-hexameric particles. The map resolution at each stage is indicated. **d,** As in (b), but for di-hexameric particles. **e,** As in (c), but for di-hexameric particles. **f,** Representative flowchart shows the computational recovery of density for the left hexamer via particle subtraction and focused 3D classification using a mask made from the density subtracted map. The particles that result in good maps are circled in red and were merged to reconstruct the hexamer map. The map resolution after masked local refinement is indicated.

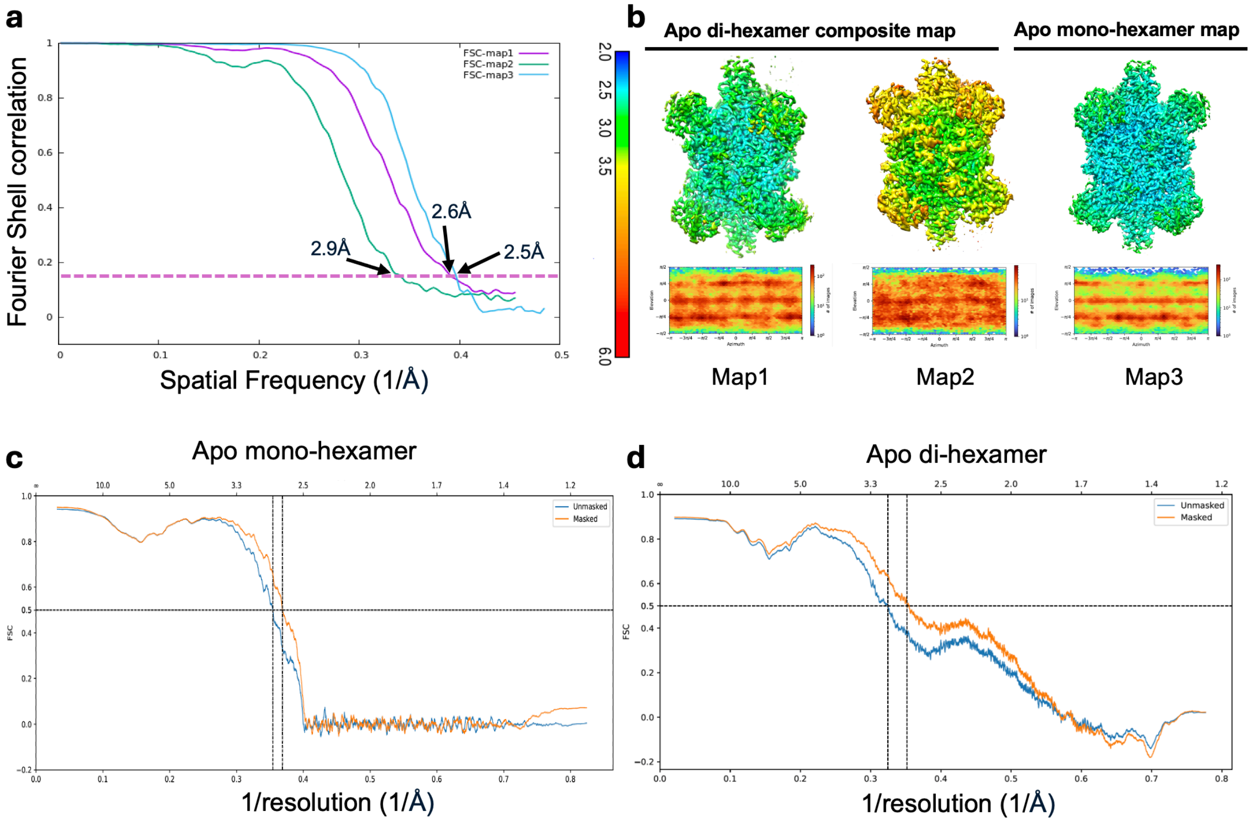

**Figure S5. Map and model validation for apo bGDH datasets. a,** Fourier shell correlation (FSC) curves for Map1-3 that was derived from half-maps with FSC cutoff 0.143 indicated, respectively, as well as nominal resolution values. **b,** Local resolution of maps (upper panel) and corresponding angular distribution plots (lower panel). Map1-3 corresponds to one hexamer from the di-hexamer, the other hexamer from the di-hexamer, and the mono-hexamer map, respectively. **c,** FSC curves derived from map-to-model reconstructions of apo mono-hexamer, with FSC cutoff 0.5 indicated. **d,** As in panel c, but for apo di-hexamer.

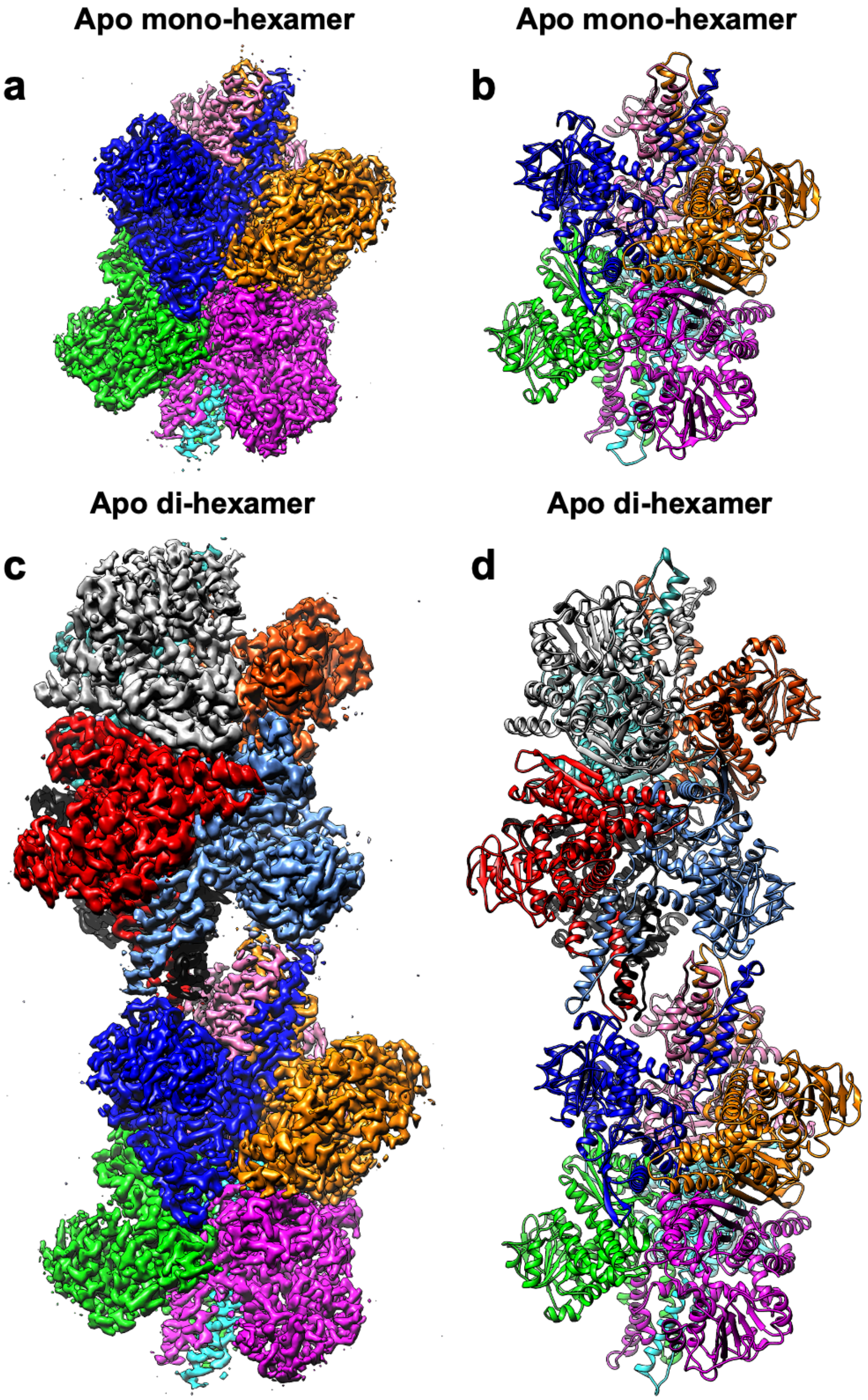

**Figure S6. Cryo-EM maps and their corresponding atomic models of bGDH mono- and di-hexamers**. **a-b**, Reconstructed cryo-EM map of apo bGDH mono-hexamer and its atomic model, respectively. **c-d**, Reconstructed cryo-EM map of apo bGDH di-hexamer and its atomic model, respectively.

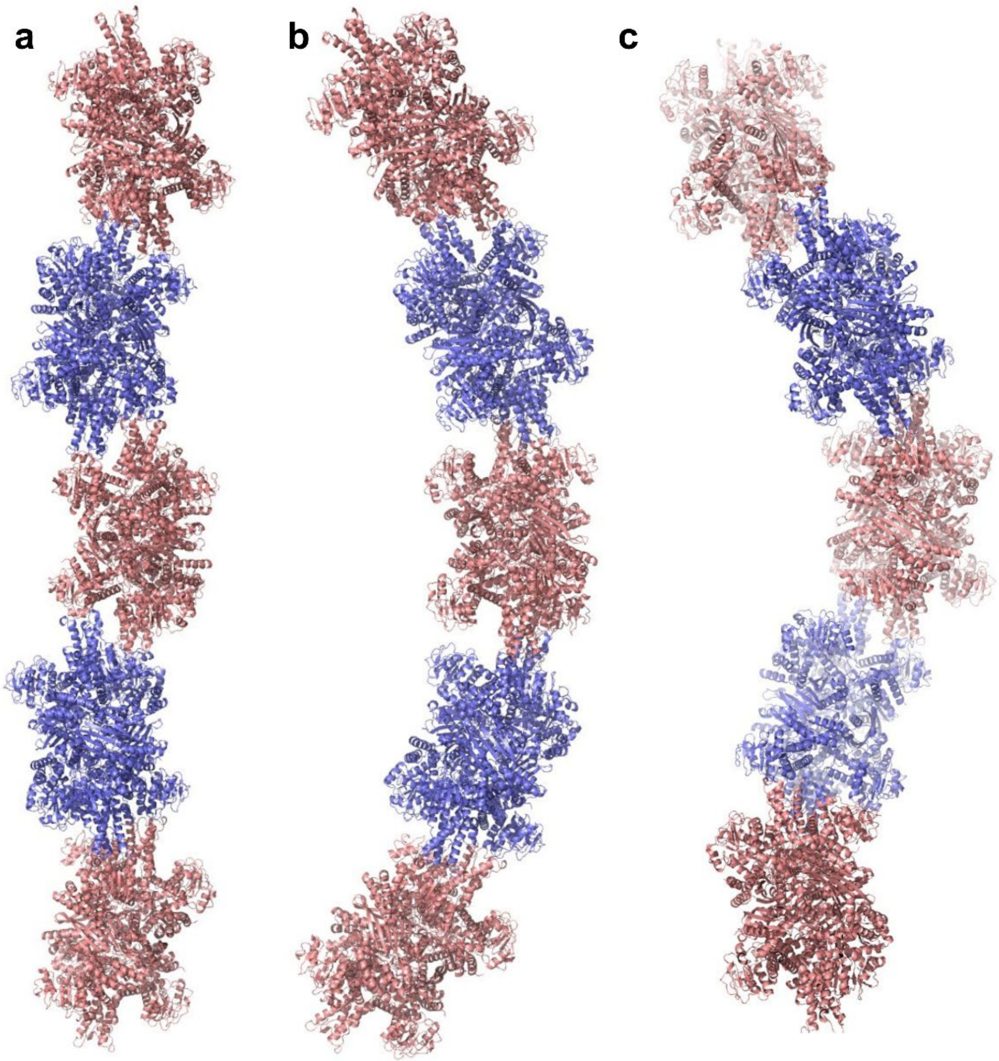

**Figure S7. Models of the bGDH filament based on the apo di-hexamer structure. a,** Due to the breaking of symmetry within the antenna at the hexamer-hexamer interface, three unique di-hexamer superpositions are possible. In this superposition, a straight filament is created. Hexamers colored in pink and blue. **b,** As in (a), using a different subunit of the antenna in the di-hexamer superposition generating a filament with gentle curvature characterized by an ~30° bend between adjacent di-hexamers. **c,** As in (a) and (b), using the third subunit of the antenna in the di-hexamer superposition generating a left-handed helical filament characterized by an ~60° bend between adjacent di-hexamers.

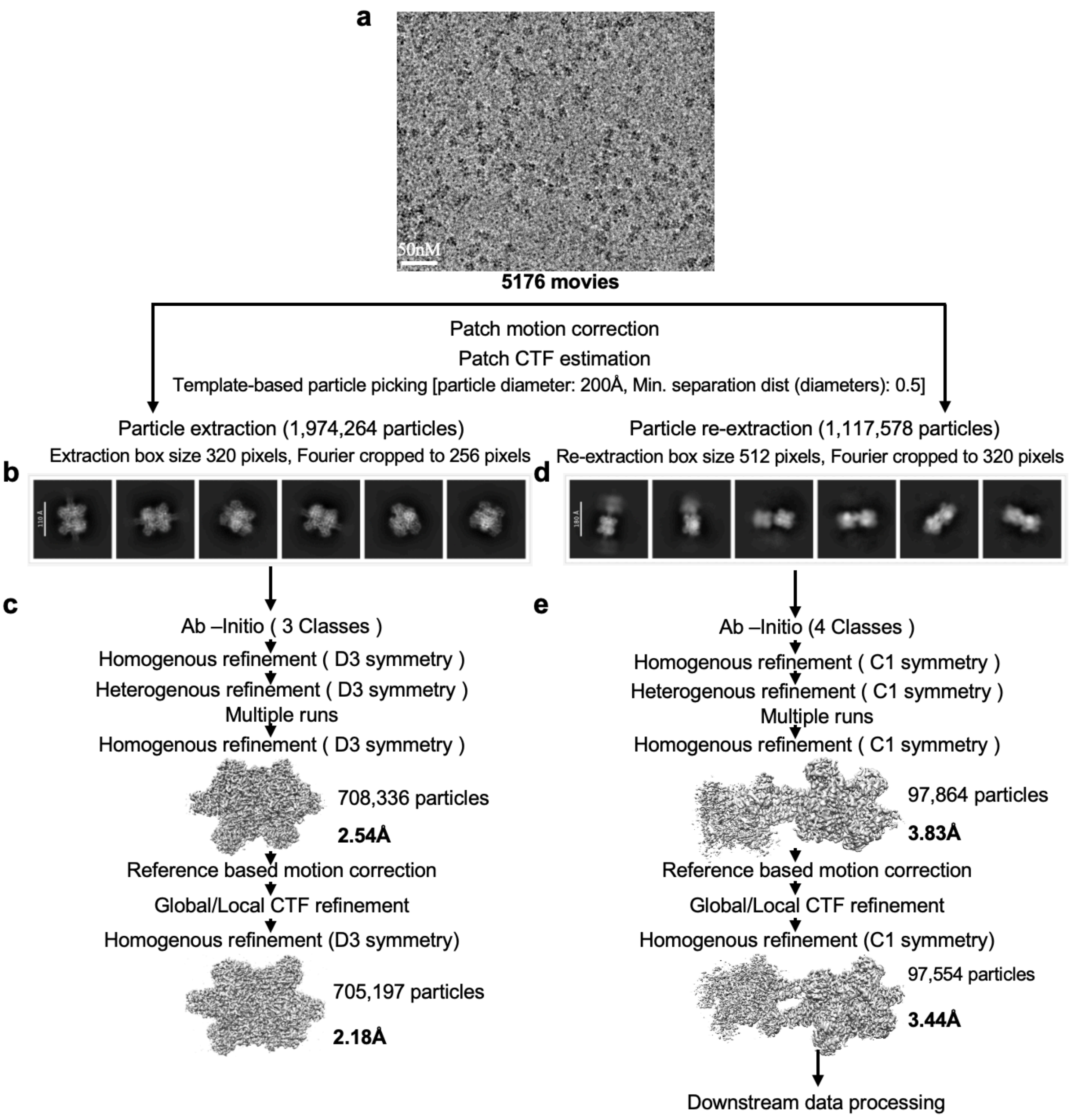
a
50nM
5176 movies
Patch motion correction
Patch CTF estimation
Template-based particle picking [particle diameter: 200Å, Min. separation dist (diameters): 0.5]
Particle extraction (1,974,264 particles)
Extraction box size 320 pixels, Fourier cropped to 256 pixels
Particle re-extraction (1,117,578 particles)
Re-extraction box size 512 pixels, Fourier cropped to 320 pixels
b
d
c
Ab –Initio ( 3 Classes )
Homogenous refinement ( D3 symmetry )
Heterogenous refinement ( D3 symmetry )
Multiple runs
Homogenous refinement ( D3 symmetry )
708,336 particles
2.54Å
Reference based motion correction
Global/Local CTF refinement
Homogenous refinement (D3 symmetry)
705,197 particles
2.18Å
e
Ab –Initio (4 Classes )
Homogenous refinement ( C1 symmetry )
Heterogenous refinement ( C1 symmetry )
Multiple runs
Homogenous refinement ( C1 symmetry )
97,864 particles
3.83Å
Reference based motion correction
Global/Local CTF refinement
Homogenous refinement (C1 symmetry)
97,554 particles
3.44Å
Downstream data processing

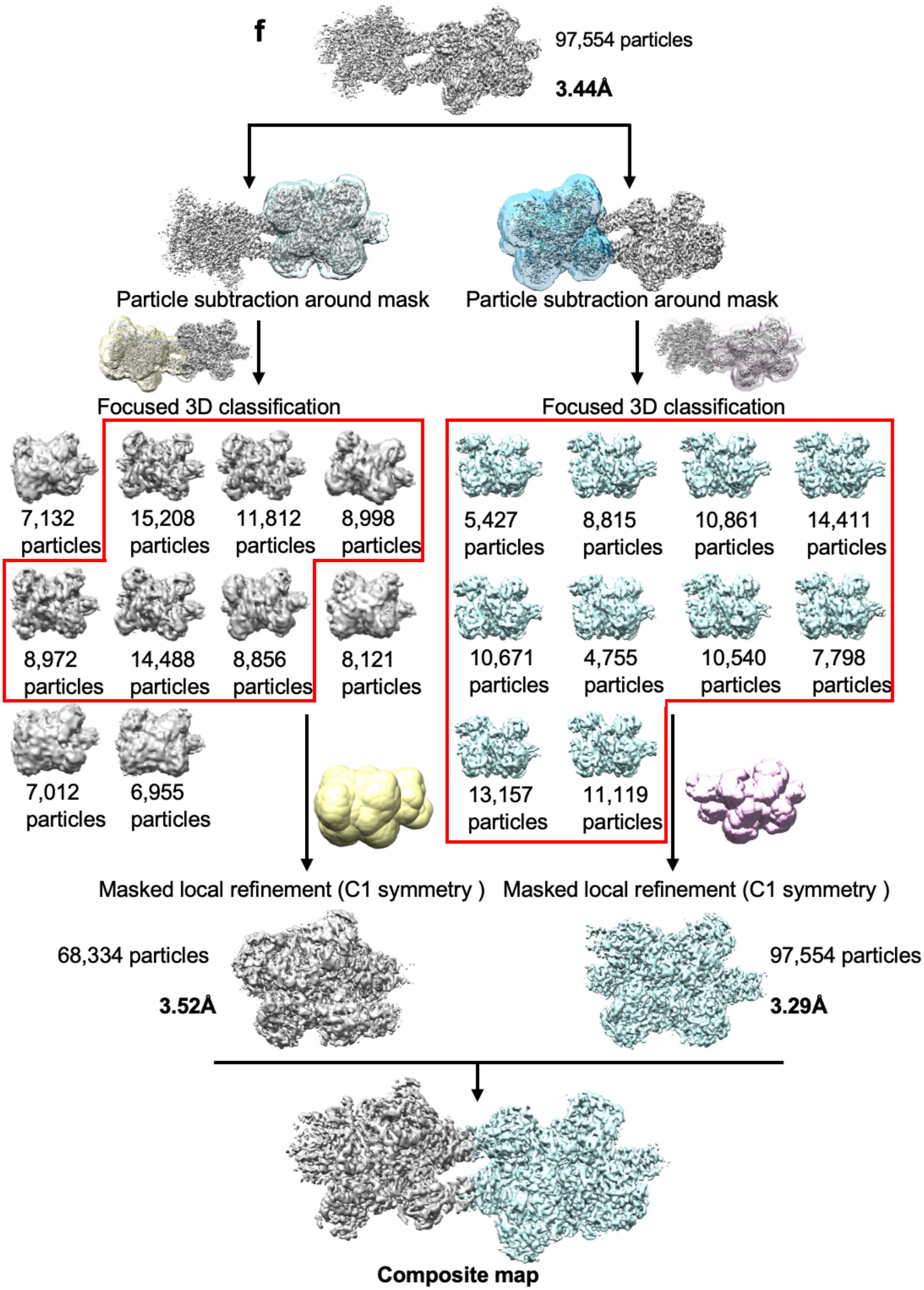
f
97,554 particles
3.44Å
Particle subtraction around mask
Particle subtraction around mask
Focused 3D classification
Focused 3D classification
7,132 particles
15,208 particles
11,812 particles
8,998 particles
5,427 particles
8,815 particles
10,861 particles
14,411 particles
8,972 particles
14,488 particles
8,856 particles
8,121 particles
10,671 particles
4,755 particles
10,540 particles
7,798 particles
7,012 particles
6,955 particles
13,157 particles
11,119 particles
Masked local refinement (C1 symmetry )
Masked local refinement (C1 symmetry )
68,334 particles
3.52Å
97,554 particles
3.29Å
Composite map

**Figure S8. Cryo-EM data collection and the initial image processing for liganded bGDH dataset. a,** Representative denoised cryo-EM micrograph, scale bar is indicated. **b,** Representative 2D class averages from reference-free 2D classification showing the mono-hexameric particles, scale bar is indicated. **c,** Representative flowchart of the initial stage of imaging processing, from 3D reconstruction to map refinement for mono-hexameric particles. The map resolution at each stage is indicated. **d,** As in panel b, but for di-hexameric particles. **e,** As in panel c, but for di-hexameric particles**. f,** Representative flowchart shows the computational recovery of density for the left hexamer via particle subtraction and focused 3D classification using a mask made from the density subtracted map. The particles that result in good maps are circled in red and were merged to reconstruct the hexamer maps. Map resolution for each hexamer after masked local refinement is indicated.

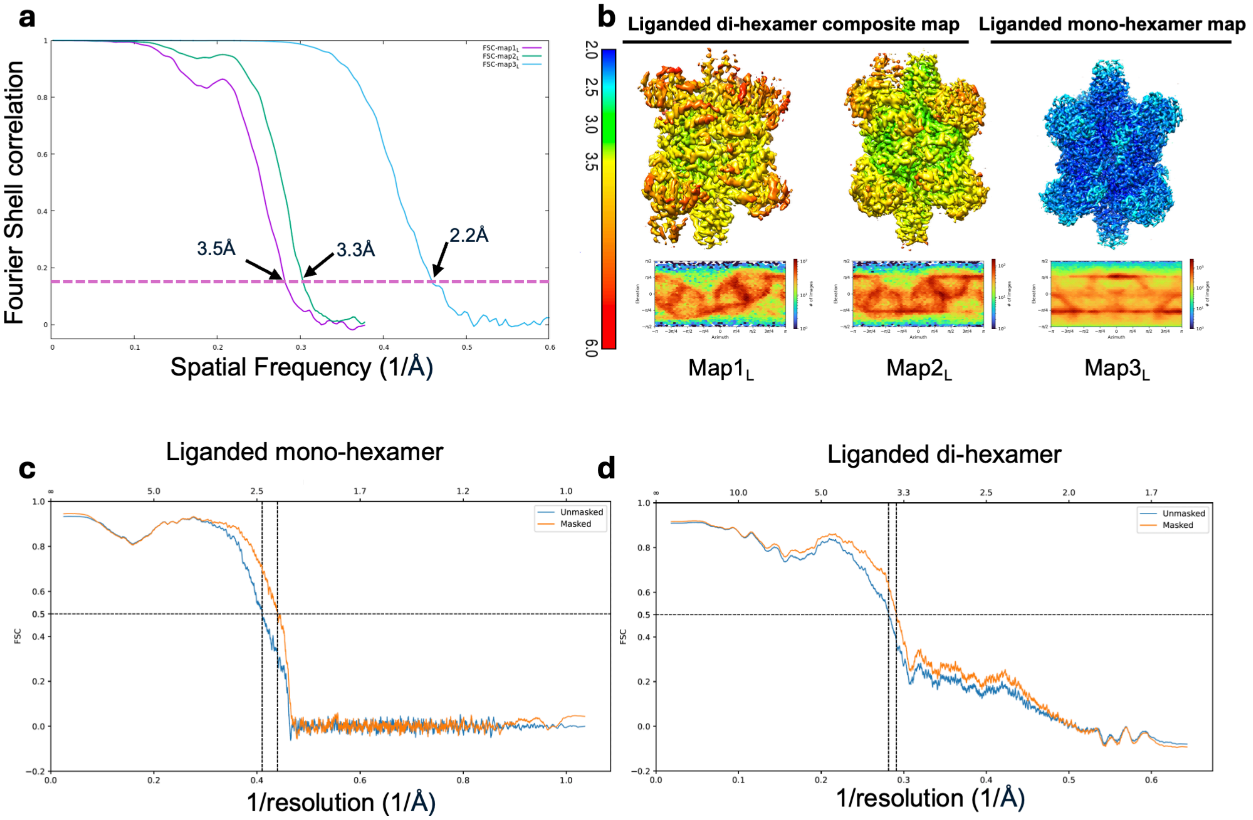

**Fig. S9. Map and model validation for liganded bGDH datasets. a,** FSC curves for Map$1_L$-$3_L$ derived from half-maps with FSC a cutoff of 0.143 indicated, respectively, as well as nominal resolution values. **b,** Local resolution of maps (upper panel) and corresponding angular distribution plots (lower panel). Map$1_L$-$3_L$ corresponds to one hexamer from the di-hexamer, the other hexamer from the di-hexamer, and the mono-hexamer map, respectively. **c,** FSC curves derived from map-to-model reconstructions of liganded mono-hexamer, with FSC cutoff 0.5 indicated. **d,** As in (c), but for the liganded di-hexamer.

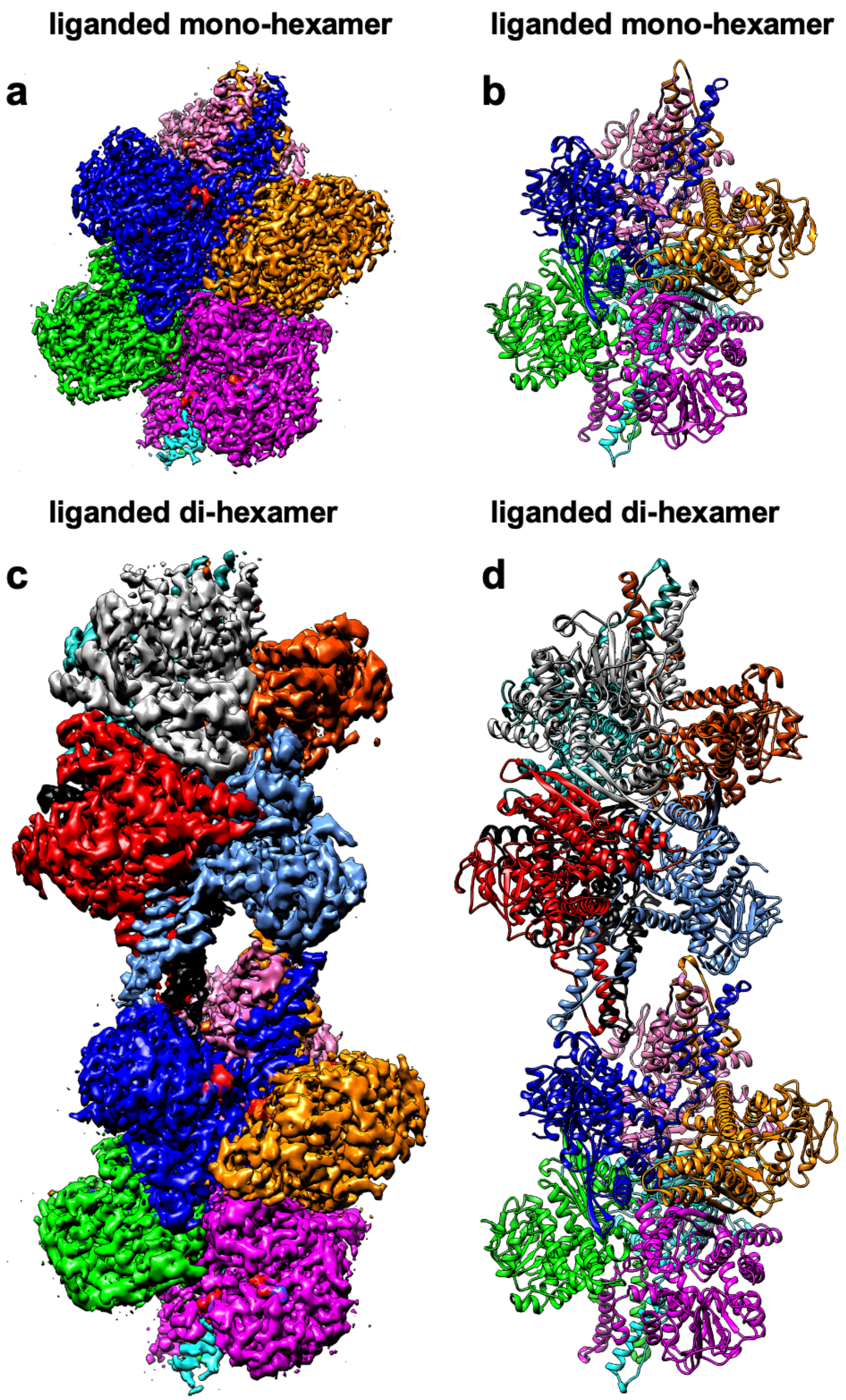

**Figure S10. Cryo-EM maps and their corresponding atomic models of bGDH mono- and di-hexamers bound to GTP, Glu, and NADH**. **a-b**, Reconstructed cryo-EM map of liganded bGDH mono-hexamer and its atomic model, respectively. **g-h,** Reconstructed cryo-EM map of liganded bGDH di-hexamer and its atomic model, respectively. The maps are colored by individual protomers.

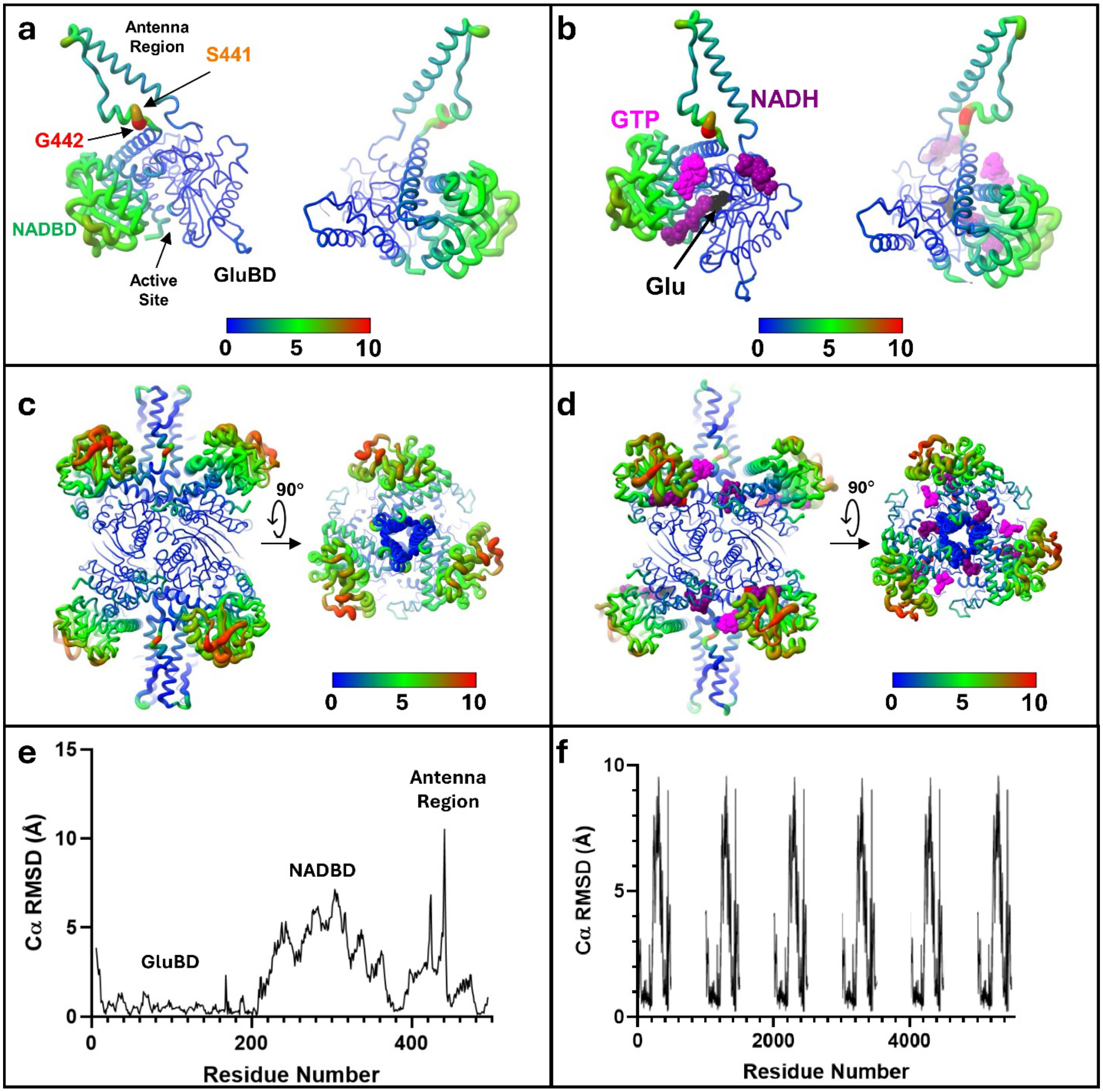

**Figure S11. RMSD analysis of ligand binding induced conformational changes in mono-hexamer structures. a,** RMSD (in Å) of Cα atoms after superposition of apo and liganded mono-hexamer structures using the glutamate binding domain (GluBD, residues 5-208, 474-496, 369-390) mapped onto a single subunit of the apo form. RMSD is plotted by color and chain thickness (thicker chain rendering indicates higher RMSD values). Larger RMSD occur in the NAD binding domain (NADBD) due to its shift to close the active site around bound ligands in the liganded structure. Large RMSDs also occur in the antenna helix and at residues Ser441 and Gly442 in the short helix following the antenna helix. **b,** As in (a), but RMSDs are mapped onto the liganded mono-hexamer subunit frame. Bound GTP, NADH, and glutamate (Glu) are shown as spheres colored pink, dark purple, and black, respectively. NADH binds in both the active site, as well as a second allosteric site. **c,** As in (a), but using all six subunits in the superposition. **d,** As in (b), but using all six subunits in the superposition. **e,** RMSD shown in (a) and (b) plotted by residue number. **f,** RMSD shown in (c)-(d) plotted by residue number. Residues numbers for different subunits are incremented by 1000 for clarity.

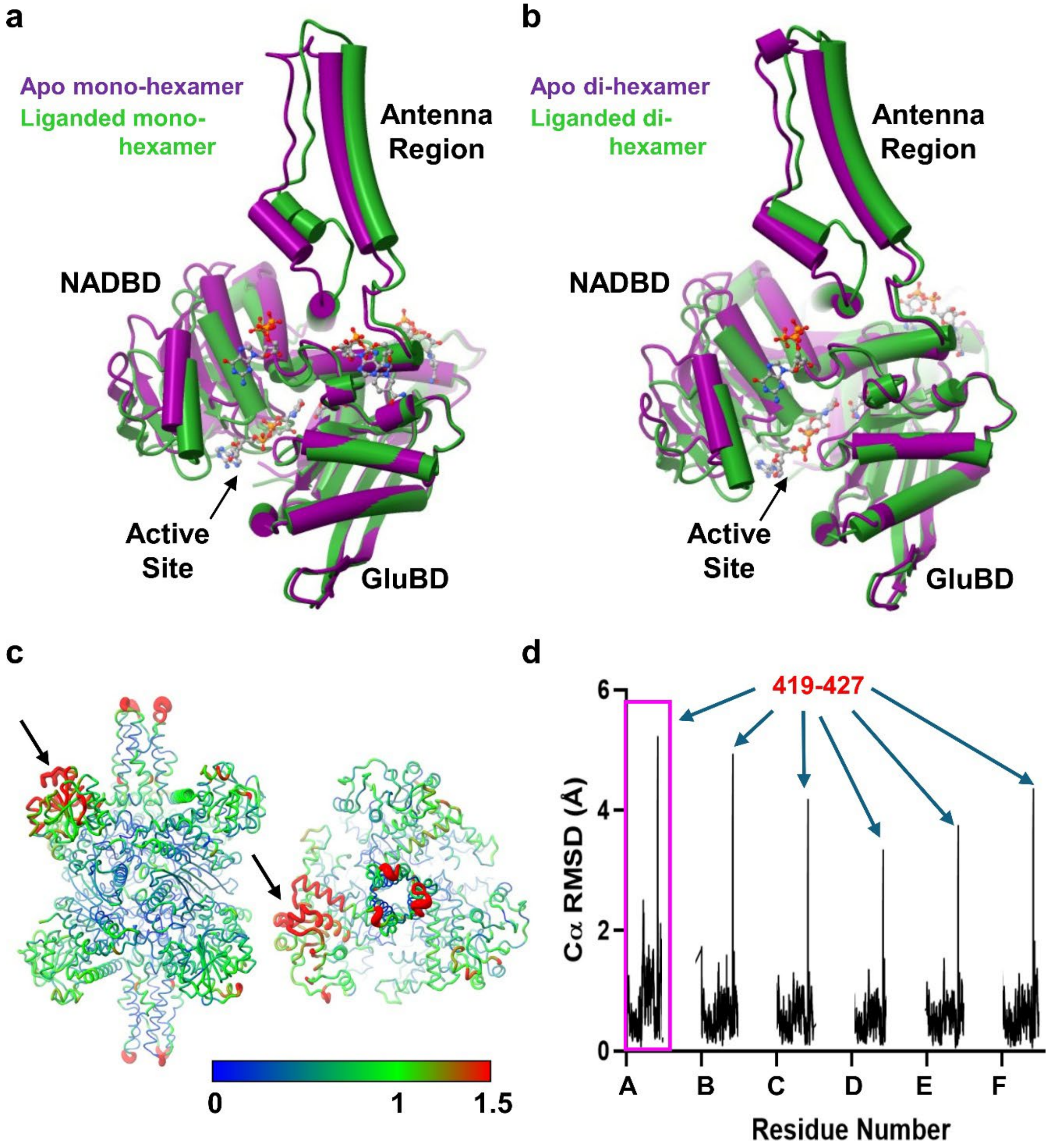

**Figure S12. Hexamer-hexamer contacts dampen conformational changes associated with ligand binding. a,** Comparison of a single subunit of bGDH from mono-hexameric structures in apo (dark purple) and liganded (green) states. The Cα atoms of the GluBD were used in the superposition. **b,** As in (a), but in the di-hexamer structures. The subunits involved in hexamer-hexamer interactions via two contact sites were used in the superposition. **c,** RMSD (in Å) of Cα atoms between mono- and di-hexamer structures both in the liganded state after superposition using the Cα atoms of all residues of all six subunits. RMSDs are indicated by both coloring and chain thickness (thicker indicates greater RMSD). Note the large RMSD at the antenna tips and in the NADBD of the subunit contacting the neighboring hexamer via its NADBD (arrow). **d,** RMSD from (c) plotted by residue number of each subunit A-F. The largest RMSDs occur at the antenna tips (residues 419-427) and within the NADBD of subunit A, which contacts the associated bGDH hexamer via its NADBD (pink box). Subunits B and C engage at the hexamer-hexamer interface via their antenna regions only.

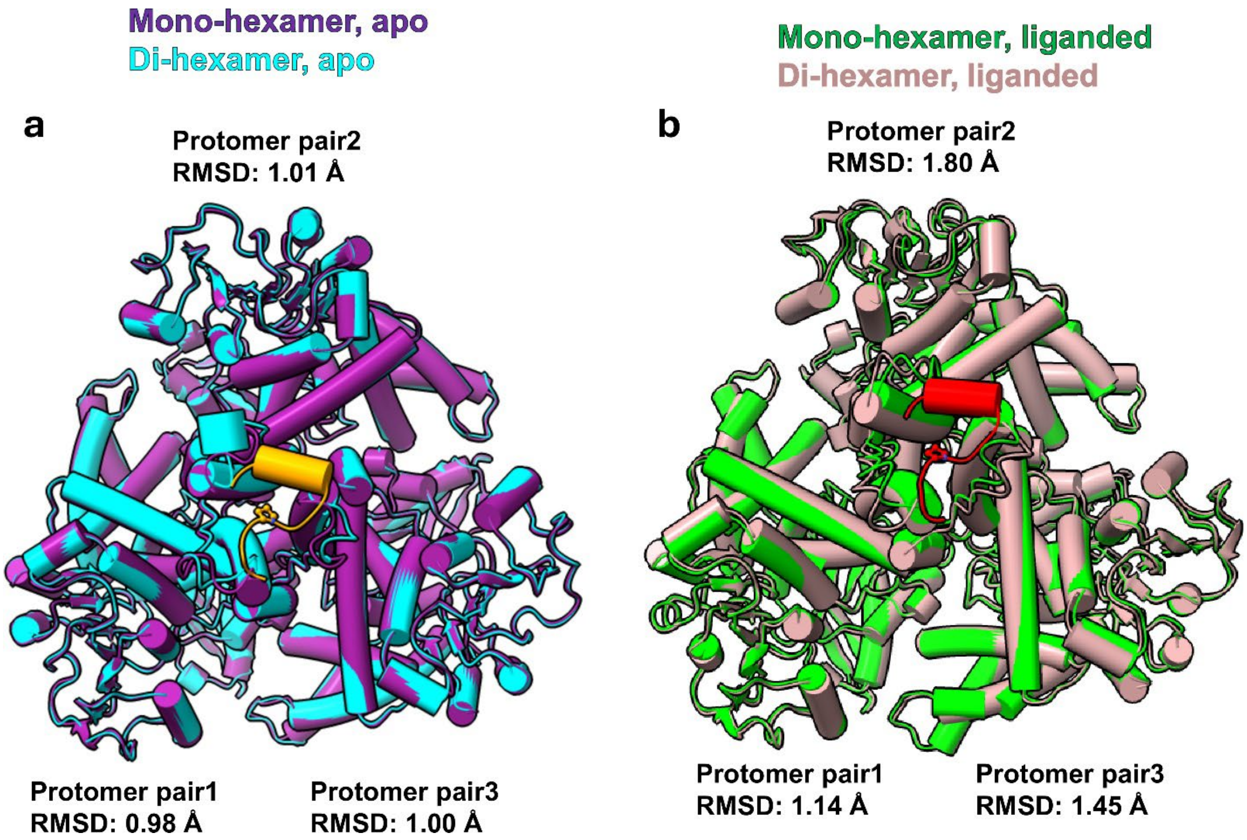

**Figure S13. RMSD between mono- and di-hexamer in apo and liganded states. a,** RMSD difference between mono- (purple) and di-hexamer (cyan) in the apo state by superimposing a single subunit found at the di-hexamer structure. The RMSD difference in each protomer pair between mono- and di-hexamer is indicated. **b,** As in (a), but using the liganded structures. The larger RMSD difference in protomer pair1 compared to the other two protomer pairs indicates that the protomer arrangement differs between liganded mono- and di-hexamer structures. The protomer with the largest RMSD (pair2) is engaged in contacts with the neighboring hexamer using both its antenna helix and its NAD binding domain.

a b

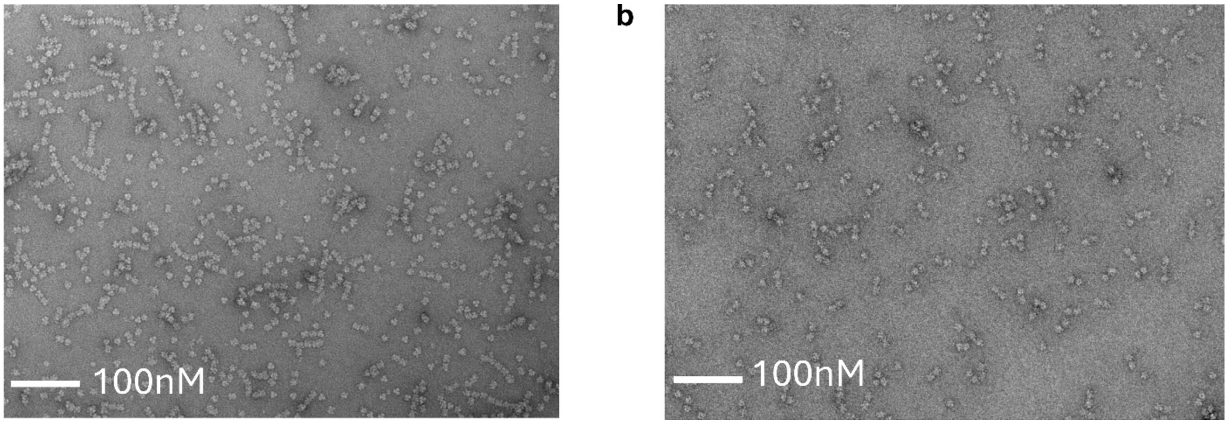

**Figure S14. Negatively stained micrographs show filaments of bGDH in assembly buffers.** **a,** bGDH (0.5 mg/ml) in a buffer containing 25 mM Tris-HCl, pH 8.0, 50 mM NaCl, 0.75 M ammonium sulfate showing the presence of filamentous assemblies. **b,** As in a, but with the addition of 5 mM glutamate, 0.4 mM NADH, and 0.9 mM GTP to the sample. The enzyme assembles into more abundant and longer filaments in the apo form than in the presence of ligands resulting in abortive complex formation. Scale bars are shown.
